# Song evolution in light of ecosystem differences: exploring effects of urbanization and ecology on temporal and frequency traits of Spotted and Eastern towhee songs

**DOI:** 10.64898/2026.05.30.728953

**Authors:** Ximena León Du’Mottuchi, Meklit Mesfin, Nicole Creanza

**Affiliations:** Department of Biological Sciences, Vanderbilt University, Nashville, TN 37212; Evolutionary Studies Initiative, Vanderbilt University, VU Box #34-1634, Nashville, TN 37235

**Keywords:** acoustic adaptation hypothesis, birdsong, computational biology, cultural evolution, ecology, sister species, song variation, urbanization

## Abstract

The Eastern towhee (*Pipilo erythrophthalmus*) and Spotted towhee (*Pipilo maculatus*) are large New World sparrows found across North America. These two species were previously classified as a single species, the Rufous-sided towhee, which was separated in 1995 based on differences in plumage, geographic range, and song. Previous studies have shown that ecological factors, such as urbanization and climate, can affect learned vocalizations, particularly frequency-related song characteristics such as minimum frequency; however, most studies have been conducted on one species in a specific location. The extensive geographic distributions of these towhee species, along with their ample publicly available song-recording data, give us the unique opportunity to assess whether ecological pressures influence song variation. Here, we extract frequency-related song features from 2916 Spotted and Eastern towhee recordings and investigate whether geography and ecology—including recording location (latitude and longitude), tree cover, urbanization (nighttime lights, human population density, and distance to road), climate zones, elevation, and ecoregions—explain patterns in these song frequency variables of this sister-species pair. Our results show that geographic location, particularly longitude, contributes more strongly to the variation of song frequency features than urbanization, environment, and climate, suggesting that culturally transmitted differences in learned song, not ecology or anthropogenic structure, drive this variation. However, there was not a clear pattern of isolation-by-distance despite the geographic patterns that we found in their songs. Further, we did not find strong support for behavioral adaptation to habitat structure, but we did find a weak signal that urbanization was associated with increased frequency in Spotted towhees. Overall, we provide a detailed study on the interactions between ecology and song evolution, and provide new insight into the evolution of birdsong.

## INTRODUCTION

Oscine songbirds use their song primarily for attracting mates and defending territories, and this song is a learned behavior that is driven by both sexual and natural selection (Catchpole and Slater 2003). Thus, variation in song characteristics, such as frequency, can reflect both evolutionary divergence and environmental adaptation and can be shaped by a combination of ecological, physiological, or cultural processes.

Efficient signal transmission in the environment is crucial for acoustic communication between the sender and receiver. Previous studies suggest that differences in birdsong, particularly in frequency and temporal structure, may be influenced by the sound transmission properties of their environment and that these song differences may be a result of adaptation towards efficient signal transmission, known as the ‘acoustic adaptation hypothesis’ (Roca et al. 2016; Morton 1975). Birdsong is prone to modification by the environment, given that different habitats impose varying acoustic effects on the signal as it travels through space. Foliage and other obstacles can interfere with song transmission, causing signal degradation and problems with song perception as a result of frequency-dependent reverberations (i.e. depending on their frequency, sounds are reflected and scattered in different ways due to the interaction of the soundwaves with obstacles in the environment, leading to sound distortion), attenuations (i.e. sound energy is weakened through spherical spreading and as it gets absorbed by air or other obstacles in the environment), and irregular amplitude fluctuations (i.e. sound intensity can vary unpredictably as it is reaching the receiver due to how soundwaves interact with the environment as they travel through space; e.g. turbulent air or temperature gradients) (Wiley 1991; Wiley and Richards 1978, 1982). In general, researchers have predicted that longer, more repetitive songs with smaller frequency bandwidths and overall lower frequencies will be more common in areas with denser vegetation, but there has been mixed support for these specific predictions (Boncoraglio and Saino 2007; Hardt and Benedict 2020; Freitas et al. 2025). Additionally, these acoustic differences may be a result of long-term evolutionary changes across generations, which can lead to cultural divergence, or may occur through short-term vocal plasticity where species adjust their songs in real time in response to the environment. For instance, analogous to humans talking in a noisy room, birds in some species experience the Lombard effect in real time, increasing the amplitude of their vocalizations in response to loud background noise (e.g. sounds of other animals, rain, urban noise) to minimize signal masking (Hardman et al. 2017). Accordingly, these changes in response to background noise suggest that songs might be performed such that the transmission of the song is maximized in the native environment.

The characteristics of birds’ local environments can potentially lead to variation in songs, either directly by affecting how sound travels or indirectly by placing selection pressures on morphological characters such as beak shape and body size. For example, songs of the white-crowned sparrow showed a negative correlation between vegetation density and both trill rates and minimum frequencies (Derryberry 2009), suggesting that aspects of this species’ song might be influenced by how sound transmits due to the structure of their habitat. Another study showed that weather parameters, including air pressure, atmospheric humidity, and air and soil temperatures seem to affect the song variability (frequency and structure parameters) of songbirds (Schäfer et al. 2017): air pressure was positively correlated with song frequency bandwidth in blue tits, humidity was positively correlated with duration of song elements in great tits, and soil temperature was positively correlated with song frequency bandwidths in blackbirds. Further, some bird species have been shown to exhibit song dialect differences along an altitudinal gradient (Nottebohm 1975), indicating that elevation may also be associated with variation in birdsong. Habitat structure and climate can also indirectly drive the variation in song properties by affecting the species morphology and physiology. A study of Darwin’s finches found that beak morphology was associated with song frequency (Huber and Podos 2006); therefore, if climate or habitat structure drives the evolution of beak morphology, it could indirectly contribute to song divergence due to the associated morphological constraints. Additionally, high temperatures have been shown to affect energy expenditure in birds, which in turn can affect song production capabilities (Coomes and Derryberry 2021).

Urban environments have properties that have been shown to further contribute to changes in birdsong. The structure of urban areas, with loud and low-frequency traffic noise, concrete ground instead of plant matter, and large obstacles, like buildings, can disrupt the acoustic signals by masking birdsong or by causing songs to echo (Slabbekoorn et al. 2007) leading to problems with sound perception. Sound frequency aspects of birdsong are therefore likely to be under different selection pressures in different contexts, especially since high-frequency sounds are more easily attenuated and degraded than low-frequency sounds, but low-frequency sounds more easily drowned out by low-frequency anthropogenic noise (Marler and Slabbekoorn 2004). Thus, in some environments, such as foliage-dense forests, lower-pitched songs might be easier to transmit to a receiver since low frequencies minimize the effects of reverberation (Ryan and Brenowitz 1985), whereas other environments, such as those with loud anthropogenic noise, may favor higher-pitched songs for reduced masking and better signal transmission. For example, studies have noted that Junco songs recorded in an urban area had higher minimum frequency than those recorded in a forest (Slabbekoorn et al. 2007). Additionally, transmission experiments of artificial sounds in the urban environment showed discrete echoes, but in the forest, they had reverberations that gradually declined in amplitude (Slabbekoorn et al. 2007), further supporting the notion that habitat structures could influence the acoustic properties of bird vocalizations in predictable directions. Altogether, these observations suggest that song divergence in Oscine birds can be affected by a combination of ecological variables, through direct as well as indirect processes that can drive both short-term and long-term evolution of song.

While it has been shown that ecology and urbanization can affect the evolution of learned songs, it remains unknown whether environmental selection pressures contribute to song differences in a widespread sister-species pair, the Eastern towhee (*Pipilo erythrophthalmus*) and Spotted towhee (*Pipilo maculatus*). Previous research showed that individual song features varied across a longitudinal gradient within each species (León Du’Mottuchi and Creanza 2025). However, the geographic variation between these two species’ songs could reflect various factors, including cultural drift, isolation by distance, and adaptation to the local environment.

By incorporating ecology and urbanization, which both have been shown to affect birdsong, we hypothesize that we can achieve more accurate song estimates and improve our understanding of song evolution in this sister species pair. Further, due to their vast combined geographic ranges expanding from the Atlantic to the Pacific ocean and from Canada to Guatemala, these species are an excellent system for understanding the evolution of learned vocalizations across a broad geographic scale and extensive range of ecological conditions. Variation in birdsong, particularly in frequency, has been previously attributed to modifications imposed by the acoustic environment. Here, we investigate whether various measures of song frequency correlate with ecosystem differences in this sister-species pair using statistical analyses of community-science recordings of birdsong. We hypothesize that bird song frequency, temporal regularity, and syntactic regularity is partly shaped by environmental conditions and will be correlated with habitat structure, degree of urbanization, and climate. This research will shed light on the potential ecological factors that may have contributed to song differences in this pair of sister species and provide insight into how learned vocalizations may evolve in response to the environment.

## METHODS

### Song Analysis

Following the methods of a previous study of towhee songs (León Du’Mottuchi and Creanza 2025), we collected song recordings of Spotted and Eastern towhees from the Macaulay Library (Cornell Lab of Ornithology 2009), a repository for bird sound recordings. We limited the recordings to those recorded during the breeding season between April and August based on the Birds of the World database (Bartos and Greenlaw 2020; Greenlaw 2020). To minimize the possibility of repeated recordings of the same bird, we categorized recordings submitted by the same recordist on the same day and in the same location as duplicates and filtered them out from our dataset. We manually selected a single bout of song from each recording, changed the sampling rate to 44100 Hz, and exported the files as .wav files (if not already in the correct format) using Audacity version 3.1.3 (https://www.audacityteam.org/) to standardize the recordings for analysis. Then, we segmented the bouts into syllables and performed song analysis using Chipper (Searfoss et al. 2020). For the song analysis, we set the noise threshold (Spotted towhee = 87, Eastern towhee = 70) and syllable similarity threshold (Spotted towhee = 26.6, Eastern towhee = 25.5) using the same parameters that were used in the previous study. We analyzed 8 different frequency metrics, including average syllable upper frequency, average syllable lower frequency, maximum syllable frequency, minimum syllable frequency, overall syllable frequency range, largest syllable frequency range, smallest syllable frequency range, and average syllable frequency range (see (Searfoss et al. 2020) for song feature calculations).

**Figure 1.**
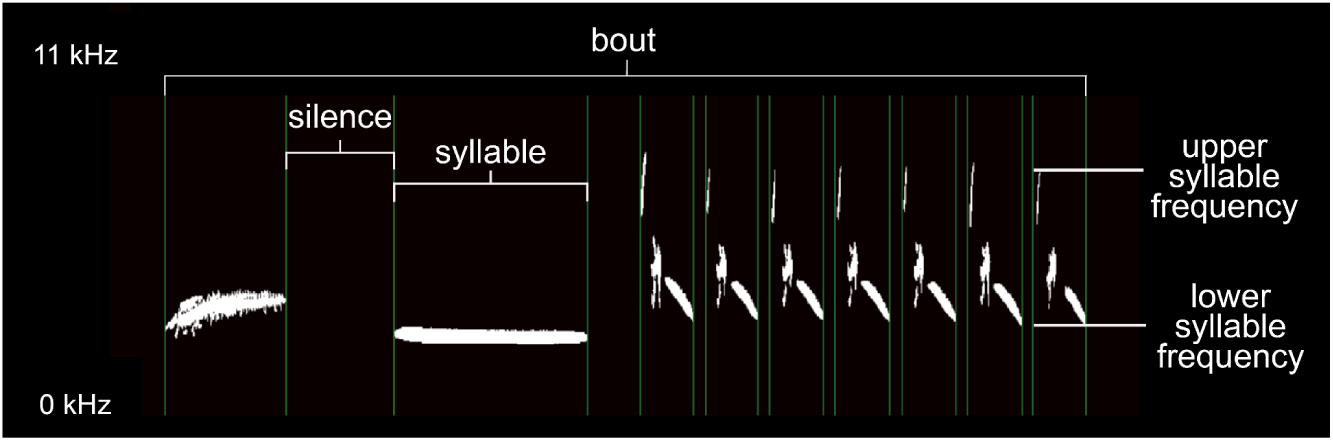
Spectrogram showing the syllable segmentation of an Eastern towhee song using the Chipper software. The spectrogram was generated from Macaulay Library recording ML564113291, https://macaulaylibrary.org/asset/564113291.

We combined these songs with the towhee song database from a previous study (León Du’Mottuchi and Creanza 2025), which also extracted song recordings from Macaulay Library as well as Xeno-canto (Planqué et al. 2005; Araya-Salas and Smith-Vidaurre 2017), bringing the total to 1830 Eastern towhee songs and 1086 Spotted towhee songs recorded between 1950 and 2024 in North America. In order to test our hypotheses with more modern data, we also separately analyzed only samples within the continental United States from 2016 through 2024 (hereafter called “modern towhee dataset”; **Fig. S1**), which could more readily be paired with recent ecology and urbanization data from 2020. This modern towhee dataset has a total of 1387 Eastern towhee recordings and 711 Spotted towhee recordings (∼72% of our total recordings) subsetted from the full towhee dataset. Since the values of the resulting song features were not normally distributed, we used the natural log of each feature in our statistical analyses to investigate whether song differences are associated with urbanization and ecology. We provide our code for statistical analysis and visualization of the song feature data and metadata about each song recording on GitHub (https://github.com/CreanzaLab/TowheeEcology).

### Environmental and Urbanization Data Collection

To observe the relationship between environmental factors and song frequency measurements, we collected data on human population density, nighttime lights, and distance to nearest road (proxies of urban noise levels), as well as canopy/tree cover, elevation, ecoregions, and climate zones (environmental characteristics). By mapping these high-resolution datasets across the species’ geographic range, we aimed to test whether the properties of a song recorded at a given location could be predicted by the environmental characteristics of that specific location.

For population density (persons/km^2^), we obtained global raster files of the estimates of the human population from the Socioeconomic Data and Applications Center (SEDAC) Gridded Population of the World dataset using the files from the year 2000 for the full towhee dataset and 2020 for the modern towhee dataset (Center for International Earth Science Information Network-CIESIN - Columbia University 2018). As another proxy of urban noise levels, we used the National Oceanic and Atmospheric Administration (NOAA) VIIRS Nighttime Lights (VNL) (Elvidge et al. 2017) stable lights (i.e. only stable, non-transient human lights included) raster data version 1 from 2013 for the full towhee dataset and version 2.1 from the year 2020 for the modern towhee dataset. We then collected tree coverage data to compare with the song characteristics; for the full towhee dataset, we used the percent tree coverage raster files from the year 2000 from Google Earth Engine (Hansen et al. 2013), and for the modern towhee dataset, we obtained raster image files of percent canopy coverage data from the year 2020 from the USDA (Housman et al. 2023). We obtained elevation raster data from the Commission for Environmental Cooperation (CEC) from 2023 (Commission for Environmental Cooperation 2023). Additionally, we obtained climate zone shapefiles for North America from the CEC based on the Köppen-Geiger climate classification (Beck et al. 2018; Peel et al. 2007)**S2 & Table S1**; (Beck et al. 2018; Peel et al. 2007). We also used the Level 1 Ecoregions of North America shapefiles from the CEC to obtain ecoregion classifications based on geographic location ((Beck et al. 2018; Peel et al. 2007)**S3 & Table S2**; (Commission for Environmental Cooperation, n.d.).

In QGIS (QGIS Development Team 2025), a geographic information system software, we imported the ecology vector and raster files as layers, along with a comma-delimited text file containing the coordinates for the towhee recordings. We then reprojected the ecoregion, climate zone, and elevation data to EPSG:4326 - WGS 84 using the ‘warp(reproject)’ tool (using the nearest neighbor method for ecoregion and climate zone files and the bilinear method for the percent canopy coverage file) for raster files and the ‘Reproject Layer’ tool for vector files to keep the layer projection consistent for the data extraction steps. With the ‘Point Sampling Tool’ in QGIS, we extracted the corresponding environmental data for the location of each towhee recording in our dataset using the longitude and latitude coordinates. If samples in our dataset were missing climate zone data or ecoregion data (i.e. when longitude/latitude coordinates of samples were rounded such that the estimated location was in the ocean), we manually assigned it to the nearest climate zone/ecoregion.

For distance to road, we used the Roads map (version 5.0.0) shapefile from Natural Earth obtained from data from the CEC North America Environmental Atlas, which shows “basic” roads at a 10-meter scale. To extract the distance to major road data we imported the roads map shapefile (EPSG:4326 - WGS 84) and the delimited text layer that contains the longitude and latitude coordinates for the song recordings into QGIS. We reprojected both layers into North America Lambert Conformal Conic (ESRI: 102009) projection to achieve more accurate measures of distance in kilometers. Then, we used the ‘Shortest line between features’ tool which calculates the straight line distance in kilometers from the coordinates of each song recording to the intersection of its nearest major road.

Finally, we generated a dataframe that includes the song feature data, geographic coordinates, urbanization and ecology data, and species classification for both the full towhee dataset and the modern towhee dataset.

### Generalized Linear Model

To assess the degree to which each environmental factor affects the variation in song frequency, we implemented a generalized linear model (GLM) for each species separately (**Table 1**). We fit the GLMs to the raw song feature data with a Gaussian family log link function using the ‘glm’ function from the stats package (R Core Team 2012) in R. Each environmental factor (longitude, latitude, population density, nighttime lights, distance to road, canopy coverage, and elevation) was scaled and incorporated as a fixed effect in the model. Additionally, we fit a GLM to the combined set of Spotted and Eastern towhee samples and used the environmental factors and species classification as fixed effects (**Table 2**). We repeated this for the full towhee dataset (**Tables S3 & S4**).

**Table 1.** Generalized linear models of ecological variables versus song frequency variables for Spotted and Eastern towhees separately in modern towhee dataset.

|  |  | Longitude |  | Latitude |  | Population Density |  | Nighttime Lights |  | Distance to Road |  | Canopy Coverage |  | Elevation |  |
| --- | --- | --- | --- | --- | --- | --- | --- | --- | --- | --- | --- | --- | --- | --- | --- |
| | | t | $p_{adj}$ | t | $p_{adj}$ | t | $p_{adj}$ | t | $p_{adj}$ | t | $p_{adj}$ | t | $p_{adj}$ | t | $p_{adj}$ |
| Average Syllable Upper Frequency | S | -3.07 | **$2.25 \times 10^{-3}$** | -2.06 | **0.04** | 1.35 | 0.18 | -0.76 | 0.45 | -2.92 | **$3.66 \times 10^{-3}$** | -1.81 | 0.07 | 3.34 | **$8.91 \times 10^{-4}$** |
|  | E | 7.1 | **$2.04 \times 10^{-12}$** | -2.81 | **$5.10 \times 10^{-3}$** | 0.11 | 0.91 | 0.77 | 0.44 | 1.16 | 0.25 | 0.74 | 0.46 | 3.27 | **$1.10 \times 10^{-3}$** |
| Average Syllable Lower Frequency | S | -0.41 | 0.68 | -1.80 | 0.07 | 0.74 | 0.46 | 0.18 | 0.86 | -2.71 | **$6.99 \times 10^{-3}$** | -1.49 | 0.14 | 0.37 | 0.71 |
|  | E | 5.27 | **$1.61 \times 10^{-7}$** | -0.29 | 0.77 | 0.11 | 0.91 | 1.82 | 0.07 | 0.15 | 0.88 | 0.62 | 0.53 | 0.57 | 0.57 |
| Maximum Syllable Frequency | S | 1.59 | 0.11 | -0.95 | 0.34 | 1.78 | 0.08 | 0.22 | 0.83 | -2.07 | **0.04** | -1.54 | 0.12 | 5.91 | **$5.48 \times 10^{-9}$** |
|  | E | 6.35 | **$2.96 \times 10^{-10}$** | -2.64 | **$8.33 \times 10^{-3}$** | 1.25 | 0.21 | -0.77 | 0.44 | 0.52 | 0.60 | -0.34 | 0.73 | 4.32 | **$1.70 \times 10^{-5}$** |
| Minimum Syllable Frequency | S | -3.01 | **$2.68 \times 10^{-3}$** | 0.71 | 0.48 | -1.16 | 0.25 | 0.25 | 0.80 | -1.01 | 0.31 | -0.59 | 0.56 | -2.49 | **0.01** |
|  | E | 1.52 | 0.13 | -5.42 | **$7.02 \times 10^{-8}$** | 0.91 | 0.36 | -0.52 | 0.61 | -1.5 | 0.13 | 0.41 | 0.68 | -0.36 | 0.72 |
| Overall Syllable Frequency Range | S | 3.12 | **$1.87 \times 10^{-3}$** | -1.35 | 0.18 | 2.29 | **0.02** | 0.07 | 0.95 | -1.29 | 0.20 | -1.17 | 0.24 | 6.86 | **$1.54 \times 10^{-11}$** |
|  | E | 5.46 | **$5.53 \times 10^{-8}$** | -0.53 | 0.59 | 0.86 | 0.39 | -0.53 | 0.59 | 1.12 | 0.26 | -0.44 | 0.66 | 4.33 | **$1.57 \times 10^{-5}$** |
| Largest Syllable Frequency Range | S | 0.13 | 0.90 | -1.53 | 0.13 | 1.19 | 0.24 | -0.47 | 0.64 | -0.45 | 0.65 | -0.38 | 0.71 | 2.82 | **$4.86 \times 10^{-3}$** |
|  | E | 2.81 | **$4.98 \times 10^{-3}$** | -2.57 | **0.01** | 0.12 | 0.91 | -0.6 | 0.55 | -0.13 | 0.9 | -0.71 | 0.48 | 2.39 | **0.02** |
| Smallest Syllable | S | -4.12 | **$4.19 \times 10^{-5}$** | 0.22 | 0.83 | -0.82 | 0.41 | $5.81 \times 10^{-4}$ | 1.00 | -0.54 | 0.59 | 0.25 | 0.80 | 1.47 | 0.14 |
| Frequency Range | E | 3.23 | **1.27x10<sup>-3</sup>** | -3.01 | **2.69x10<sup>-3</sup>** | -0.7 | 0.49 | 0.05 | 0.96 | 0.83 | 0.40 | -0.42 | 0.67 | 0.46 | 0.65 |
| Average Syllable Frequency Range | S | -2.51 | **0.01** | -0.42 | 0.67 | 0.71 | 0.48 | -0.84 | 0.40 | -0.48 | 0.63 | -0.48 | 0.63 | 2.85 | **4.50x10<sup>-3</sup>** |
|  | E | 3.32 | **9.32x10<sup>-4</sup>** | -2.63 | **8.54x10<sup>-3</sup>** | 5.00x10<sup>-3</sup> | 1.00 | -0.50 | 0.62 | 1.02 | 0.31 | 0.34 | 0.74 | 2.85 | **4.47x10<sup>-3</sup>** |

**Table 2.** Generalized linear model of ecological variables versus song frequency variables in modern towhee dataset.

|  | Species |  | Longitude |  | Latitude |  | Population Density |  | Nighttime Lights |  | Distance to Road |  | Canopy Coverage |  | Elevation |  |
| --- | --- | --- | --- | --- | --- | --- | --- | --- | --- | --- | --- | --- | --- | --- | --- | --- |
| | t | $p_{\text{adj}}$ | t | $p_{\text{adj}}$ | t | $p_{\text{adj}}$ | t | $p_{\text{adj}}$ | t | $p_{\text{adj}}$ | t | $p_{\text{adj}}$ | t | $p_{\text{adj}}$ | t | $p_{\text{adj}}$ |
| Average Syllable Upper Frequency | 11.21 | **2.33x10<sup>-28</sup>** | 3.36 | **7.84x10<sup>-4</sup>** | -2.09 | **0.04** | 1.27 | 0.21 | -0.59 | 0.56 | -1.95 | 0.05 | 1.16 | 0.24 | 0.76 | 0.45 |
| Average Syllable Lower Frequency | 6.34 | **2.89x10<sup>-10</sup>** | 4.29 | **1.83x10<sup>-5</sup>** | -0.99 | 0.32 | 0.7 | 0.48 | 1.14 | 0.26 | -2.52 | **0.01** | 0.38 | 0.70 | -1.06 | 0.29 |
| Maximum Syllable Frequency | 6.67 | **3.26x10<sup>-11</sup>** | 6.26 | **4.63x10<sup>-10</sup>** | -1.79 | 0.07 | 2.05 | **0.04** | -0.69 | 0.49 | -1.64 | 0.10 | -0.56 | 0.58 | 6.36 | **2.47x10<sup>-10</sup>** |
| Minimum Syllable Frequency | 6.8 | **1.39x10<sup>-11</sup>** | -1.81 | 0.07 | -2.54 | **0.01** | -0.34 | 0.73 | -0.11 | 0.91 | -1.77 | 0.08 | 0.72 | 0.47 | -6.87 | **8.60x10<sup>-12</sup>** |
| Overall Syllable Frequency Range | 3.25 | **1.16x10<sup>-3</sup>** | 6.8 | **1.39x10<sup>-11</sup>** | -0.64 | 0.52 | 2.04 | **0.04** | -0.59 | 0.55 | -0.7 | 0.48 | -0.86 | 0.39 | 9.17 | **1.16x10<sup>-19</sup>** |
| Largest Syllable Frequency Range | 4.3 | **1.79x10<sup>-5</sup>** | 2.02 | **0.04** | -2.49 | **0.01** | 1.00 | 0.32 | -0.96 | 0.34 | -0.48 | 0.63 | -0.52 | 0.6 | 2.89 | **3.94x10<sup>-3</sup>** |
| Smallest | 5.69 | **1.49x10<sup>-8</sup>** | -3.62 | **3.05x10<sup>-4</sup>** | -0.22 | 0.83 | -1.15 | 0.25 | -0.20 | 0.84 | -0.27 | 0.79 | 1.32 | 0.19 | -0.76 | 0.45 |

| Syllable Frequency Range |  |  |  |  |  |  |  |  |  |  |  |  |  |  |  |  |
| --- | --- | --- | --- | --- | --- | --- | --- | --- | --- | --- | --- | --- | --- | --- | --- | --- |
| Average Syllable Frequency Range | 5.79 | **8.25x10<sup>-9</sup>** | -0.16 | 0.87 | -1.12 | 0.26 | 0.75 | 0.46 | -1.35 | 0.18 | -0.04 | 0.96 | 0.84 | 0.40 | 1.71 | 0.09 |

A generalized linear model was applied to raw song feature data of the Eastern Towhee (*Pipilo erythrophthalmus*) [ $N_{\text{Eastern}} = 1358$ ] and the Spotted Towhee (*Pipilo maculatus*) [ $N_{\text{Spotted}} = 710$ ] using a Gaussian family log link function with scaled ecological variables. After multiple hypothesis correction with FDR, results with $p_{\text{adj}} < 0.05$ are bolded and highlighted in red.

To visualize the relationship between each environmental factor and the song features, we plotted each log-transformed song feature against the longitude and latitude coordinates of its recording, population density, nighttime lights, distance to road, canopy coverage, and elevation for the modern towhee dataset (**Figs. S4-S10)**.

### Ecoregions and Climate Divisions

To assess whether song frequency variables were different in each ecoregion or climate zone relative to the overall species distribution, we used a set of Wilcoxon rank-sum tests. We used the ‘wilcox.test’ function and the ‘p.adjust’ function from the stats package in R to apply a Wilcoxon rank-sum test for each comparison with the False Discovery Rate (FDR) correction for multiple comparisons. For each species, we removed ecoregions or climate zones from the analysis if they contained fewer than 10 samples. Thus, we analyzed a total of 8 climate zones and 7 ecoregions for the Spotted towhees and 3 climate zones and 3 ecoregions for the Eastern towhees in the modern towhee dataset. In the full towhee dataset, we analyzed 10 climate zones and 8 ecoregions for the Spotted towhees and 4 climate zones and 4 ecoregions in the Eastern towhees. Additionally, we made violin plots showing the distribution of song frequency traits in each climate zone and ecoregion for the Eastern and Spotted towhee (**Figs. S11-S18**).

### Geographic Distance

To analyze whether geographic distance contributes to the difference in songs of the Spotted and Eastern towhees, we constructed a pairwise geographic distance matrix as well as a pairwise song-feature distance matrix for each combination of song recordings. We approached the song-feature distance in two ways: 1) we combined the 8 song features into a single multivariate Euclidean distance matrix to assess whether overall song similarity correlates with geographic distance (*Song Distance Model*), and 2) we analyzed the Euclidean distance of each song feature separately to assess if a specific feature correlates with geographic distance (*Song Feature Distance Models*). For both models, we applied a z-score transformation (subtracting the mean and dividing by the standard deviation) to the log-transformed song feature data to standardize the song features to a mean of zero and standard deviation of one. For the geographic distance matrix, we used the ‘geodist’ function from the geodist package (Padgham and Sumner 2025) in R to calculate the geodesic distances in meters for each pair of samples using the longitude and latitude coordinates. We then followed with the ‘as.dist’ function from the stats package to convert the matrix to a distance object. We applied the ‘dist’ function with the Euclidean distance measure from the stats package to the song feature dataframe to compute the distance matrices for the standardized song features. For the *Song Distance Model*, we applied this function to the data frame containing all 8 song features together, and for the *Song Feature Distance Models*, we applied this to a single song feature column. To identify whether there is a correlation between the geographic distance and song feature distance matrices, we used the ‘mantel’ function from the vegan package (Oksanen et al. 2024) to apply a Mantel test (Mantel 1967) based on Pearson’s product-moment correlation with 999 permutations and calculate the Mantel statistic (r) and the *p*-values. For the *Song Feature Distance Models*, we used FDR corrected *p*-values for multiple comparisons. We repeated all analyses for the Spotted towhee and Eastern towhee separately.

Additionally, to test if there is evidence of isolation by distance within each ecoregion, we used a Mantel test to calculate the *r* statistic for a given song feature in each ecoregion with FDR correction for multiple comparisons. We did this by making a song feature and geographic distance matrix for each ecoregion and song feature pair separately, and calculated the *r* statistic and FDR adjusted *p*-values using the same methods as described above.

We also assessed whether there was a stronger geographic pattern within ecoregions (i.e. a stronger association between song distance and geographic distance) than we would expect given the distribution of the species as a whole. To do this, we generated a species-wide null distribution of the Mantel *r* statistic by randomly sampling the same number of recordings (i.e. the number found in the ecoregion) from across the entire species range and computing the Mantel test statistic (using the setting permutation = 0) between their geographic and song feature distances; we then repeated this resampling procedure 999 times. We then obtained an empirical *p*-value by calculating the proportion of null *r* values that were greater than or equal to the observed *r* values for each song feature for each ecoregion. We adjusted the *p*-value for multiple comparisons using the FDR method. This allowed us to determine if within-region geographic structure in song features was stronger than what was expected by chance based on the species-wide structure.

Finally, we tested whether there is a difference in song features between ecoregions when accounting for geographic distance. For each species, we computed geographic distance versus song distance matrices as described above for each song feature. We then fit a linear regression model and extracted the residuals to remove the effects of geographic distance using the ‘lm’ and ‘resid’ functions in R. Using the ‘t.test’ function in R, we performed a t-test to assess whether the mean residual dissimilarity between each pair of ecoregions differed significantly from the average residual dissimilarity within ecoregions. We calculated the t-statistic and FDR corrected *p*-values for multiple comparisons.

These methods were applied to both the modern towhee dataset and the full towhee dataset.

## RESULTS

### Generalized linear model

We used a GLM to analyze the tree cover, nighttime lights, population density, distance to nearest road, elevation, and species data and their correlation with various frequency variables, including average syllable upper and lower frequency, maximum and minimum syllable frequency, overall syllable frequency range, largest and smallest syllable frequency range, and the average syllable frequency range. We also plotted each song feature against each environmental feature (i.e. canopy coverage, nighttime lights, population density, distance to road, elevation) for each sample in our analysis to visualize the trends (**Figs. 4 & S4-S10**).

**Figure 4.**
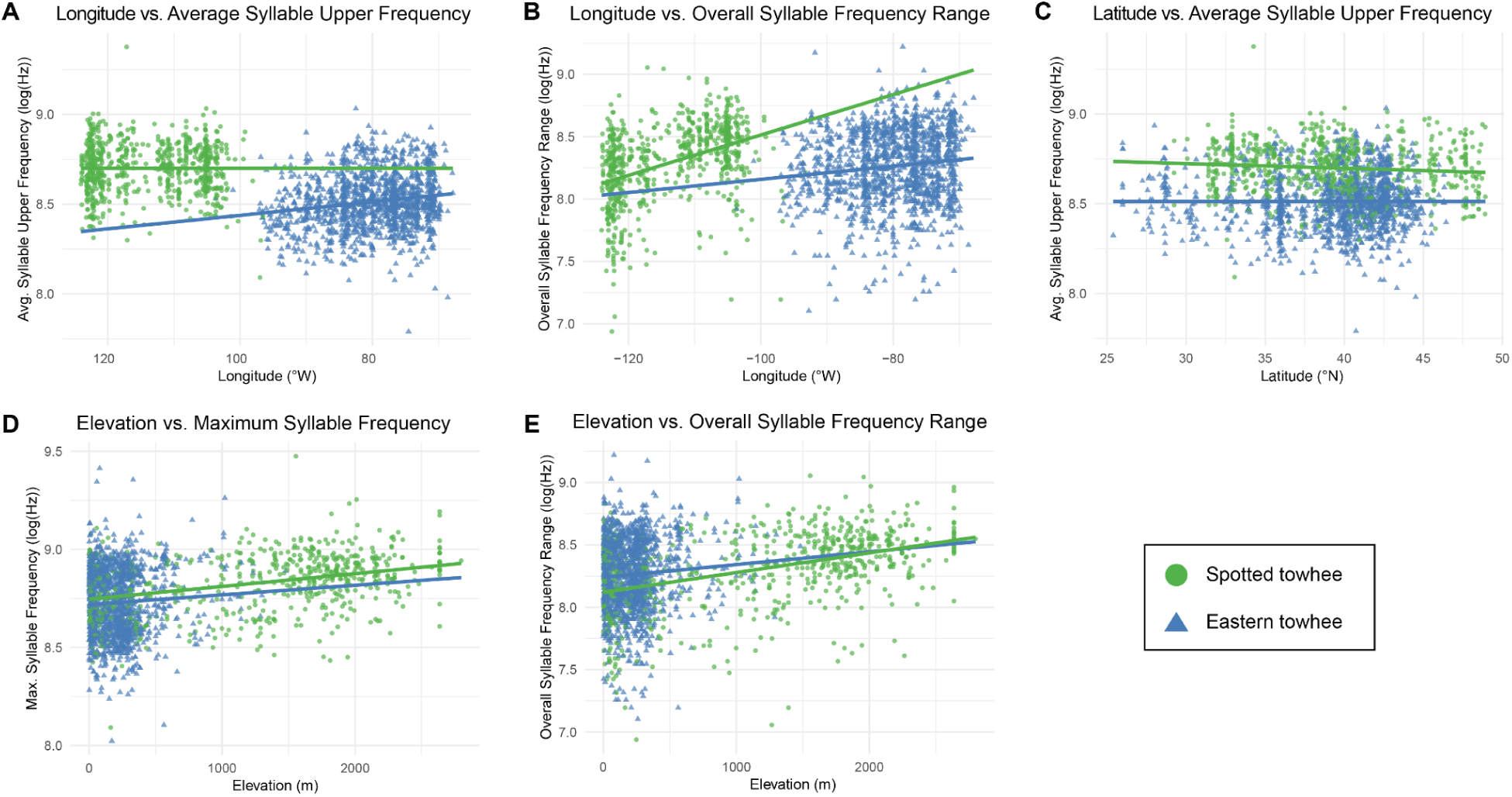
Scatterplots showing the relationship between log-transformed song frequency features (log(Hz)) and (**A–B**) longitude, (**C**) latitude, and (**D–E**) elevation. Each point represents an analyzed song in Eastern towhees (blue triangles) and Spotted towhees (green circles). A linear model was fit to the data from each species separately (indicated by lines in the corresponding color). (**A–C**) N_Eastern_towhee_ = 1387 and N_Spotted_towhee_ = 711 and (**D–E**) N_Eastern_towhee_ = 1358 and N_Spotted_towhee_ = 710; see **Table 1** for *t* and *p*-values from the generalized linear model.

For the Spotted towhee, all environmental variables except for nighttime lights and canopy coverage had at least one significant relationship (**Table 1**). Longitude was the most significant predictor of song frequency features, showing a statistically significant negative relationship with average syllable upper frequency, minimum syllable frequency, smallest syllable frequency range, and average syllable frequency range, and a significant positive relationship with overall syllable frequency range (**Fig. S4**). Latitude has a significant negative relationship with average syllable upper frequency (**Fig. S5**). Population density has a significant positive relationship with overall syllable frequency range (**Fig. S6**). Distance to road has a statistically significant negative correlation with average syllable upper frequency, average syllable lower frequency, and maximum syllable frequency (**Fig. S8**). Elevation has positive significant relationships with average syllable upper frequency, maximum syllable frequency, overall syllable frequency range, largest syllable frequency range, and average syllable frequency range, and a significant negative relationship with minimum syllable frequency (**Fig. S10**).

For Eastern towhee, longitude, latitude, and elevation were the only environmental variables that had significant relationships with song frequency variables, with the most significant predictor being longitude (**Table 1**). Again, longitude was a significant predictor (*p*_adj_ < 0.05) of 7 song features (average syllable upper frequency, average syllable lower frequency, maximum syllable frequency, overall syllable frequency range, largest syllable frequency range, smallest syllable frequency range, and average syllable frequency range), showing strong positive relationships with each feature (**Fig. S4**). Latitude showed a strong significant negative relationship with 6 song features (average syllable upper frequency, maximum syllable frequency, minimum syllable frequency, largest syllable frequency range, smallest syllable frequency range, and average syllable frequency range; **Fig. S5**). Elevation showed significant positive relationships with average syllable upper frequency, maximum syllable frequency, overall syllable frequency range, largest syllable frequency range, and average syllable frequency range (**Fig. S10**).

Further, the combined Spotted and Eastern towhee GLM model showed species classification as contributing the most to the variation in song frequency variables with statistical significance (*p* < 0.05) for all 8 features (**Table 2**). Similar to the Eastern towhee and Spotted towhee models, the second largest contributor was longitude, showing strong statistical significance in 6 song features. The GLM analyses using the full towhee dataset showed similar results (**Tables S3 & S4**).

A set of generalized linear models was applied to raw song feature data of the Eastern towhee (*Pipilo erythrophthalmus*) [N_Eastern_ = 1358] and the Spotted towhee (*Pipilo maculatus*) [N_Spotted_ = 710] using a Gaussian family log link function with scaled ecological variables. Each song feature was tested separately as a target variable, in the general format “Song Feature ∼ Longitude + Latitude + Population Density + Nighttime Lights + Distance to Road + Canopy Coverage + Elevation”. Adjusted *p*-values are shown; results with *p*_adj_ < 0.05 are bolded and highlighted in red.

A generalized linear model was applied to raw song feature data of the Eastern Towhee (*Pipilo erythrophthalmus*) [N_Eastern_ = 1358] and the Spotted Towhee (*Pipilo maculatus*) [N_Spotted_ = 710] using a Gaussian family log link function with scaled ecological variables. After multiple hypothesis correction with FDR, results with *p_adj_* < 0.05 are bolded and highlighted in red.

### Climate Zones

We applied a Wilcoxon rank-sum test to the climate zones to assess whether there was a significant difference between the song frequency variables of each species within a climate zone in comparison with the overall distribution of that species. Similarly, we found that the Spotted towhees have more differences in their frequency traits relative to the overall species distribution than the Eastern towhees in both the modern towhee dataset (**Figs. S11 & S12**) and the full towhee dataset (**Figs. S13 & S14**).

For the Spotted towhee, average syllable upper frequency, smallest syllable frequency range, and average syllable frequency range each showed a significant difference (*p*_adj_ < 0.05) in song traits compared to the overall distribution only in the humid continental dry cool summer climate zone. Largest syllable frequency range was significant for humid continental dry cool summer and warm summer Mediterranean climate zones. Maximum syllable frequency was significant for several climate zones: humid continental mild summer wet all year, warm summer Mediterranean, and cold semi-arid. Minimum syllable frequency and overall syllable frequency range were significant for warm summer Mediterranean, hot summer Mediterranean, and cold semi-arid climate zones. Overall syllable frequency range also showed significance for the climate zone humid continental mild summer wet all year, while average syllable lower frequency was not significant for any climate zone.

The Eastern towhee showed significant song differences for one climate zone compared to the overall distribution: the humid continental mild summer wet all year climate zone differed from the overall distribution for average syllable upper frequency, maximum syllable frequency, and overall syllable frequency range.

### Ecoregions

We used a set of Wilcoxon rank-sum tests to analyze whether there were significant differences between the song-frequency variables of each species within an ecoregion in comparison with the overall distribution of that species. In the modern towhee dataset, we found that both the Spotted towhees and the Eastern towhees showed a significant difference (*p*_adj_ < 0.05) in the average syllable upper frequency, average syllable lower frequency, maximum syllable frequency, overall syllable frequency range, and smallest syllable frequency range only in the Great Plains compared to the overall species-specific distribution across ecoregions (**Figs. S15 & S16**). However, in the full towhee dataset, Spotted towhees had more differences in their frequency traits within ecoregions when compared to the overall species distribution than did the Eastern towhees (**Figs. S17 & S18**).

### Geographic Distance

We used a Mantel test (Mantel 1967) to assess whether there was an association between the geographic distance matrix and each of the song feature matrices; such an association would suggest that songs accumulate changes in a pattern consistent with isolation-by-distance. The results for the Mantel tests comparing the geographic distance matrix with the multivariate distance matrices showed that geographic distance and song feature distance tend to increase together, but the relationship was weak (r < | 0.17 |, *p*_adj_ < 0.05) for the combined Spotted and Eastern towhees and the analysis of only the Spotted towhee in both the modern towhee dataset and the full towhee dataset (**Table S5**).

When analyzing the comparison between geographic distance and each song feature separately, the results showed statistical significance (r < | 0.22 |, *p*_adj_ < 0.05) in 6 song features in the combined Spotted and Eastern towhee analysis and in the analysis of only the Spotted towhee, all with a weak positive relationship for both the modern towhee dataset and the full towhee dataset (**Tables S6 and S7**). In the analysis of only the Eastern towhee in the full towhee dataset, only average syllable upper frequency was significant with a very weak positive relationship (r = 0.032; *p*_adj_ = 0.016).

### Geographic Distance with Ecoregions

Further, we tested whether there was evidence that geographic distance affects distance in song features within each ecoregion for Spotted and Eastern towhees separately. In the modern towhee dataset, we found that geographic distance was predictive of average syllable frequency range, largest syllable frequency range, maximum syllable frequency, overall syllable frequency range, and smallest syllable frequency range in the Spotted towhees only in the Northwestern Forested Mountains, but they all showed a weak positive correlation (r < | 0.27 |, *p*_adj_ < 0.05; **Fig. 5**). In the full towhee dataset, geographic distance also showed a statistically significant weak correlation with maximum syllable frequency (*r* = 0.18; *p*_adj_ = 0.03) and overall syllable frequency range (*r* = 0.23; *p*_adj_ = 0.03) in the Spotted towhees in the Northwestern Forested Mountains, in addition to showing a very weak correlation with average syllable upper frequency in Eastern towhees in Eastern Temperate Forests (*r* = 0.03; *p*_adj_ = 0.03; **Fig. S19**).

**Figure 5.**
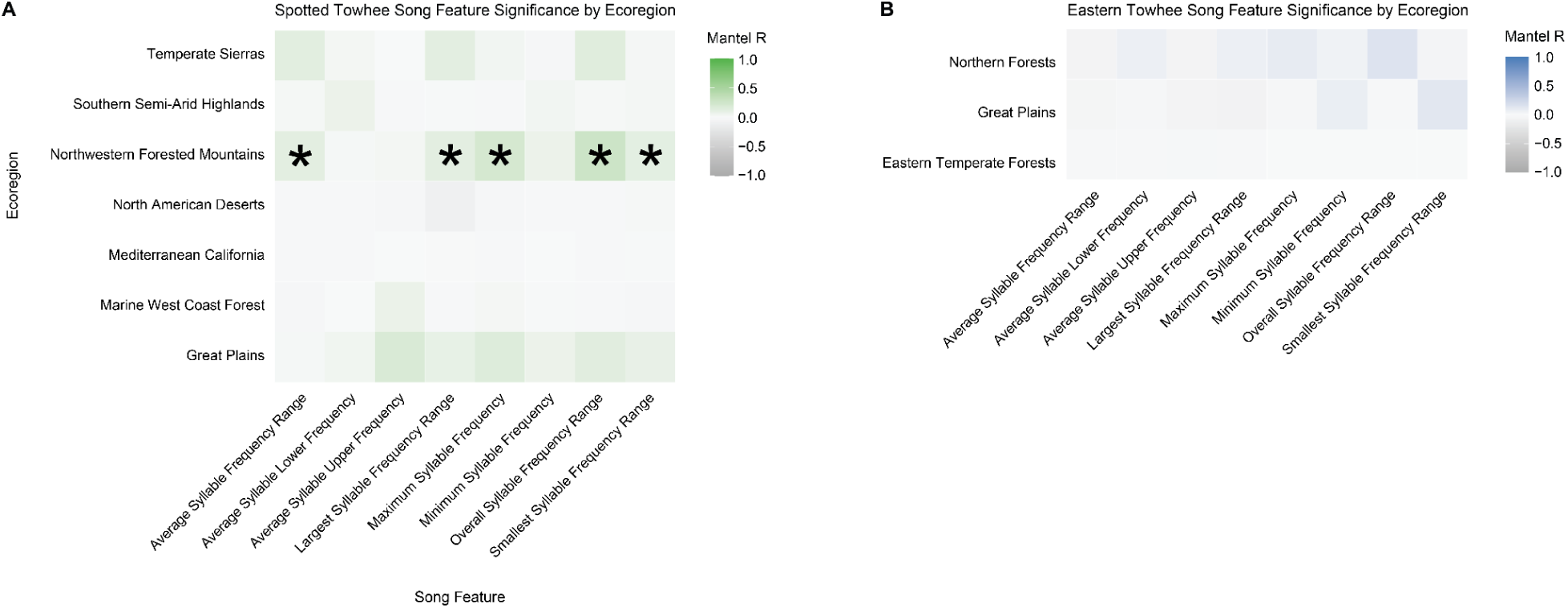
Heat map of Mantel test testing for evidence for isolation by distance for 8 song features within each ecoregion for songs of the Spotted towhee (*Pipilo maculatus*) and Eastern towhee (*Pipilo erythrophthalmus*) in the modern towhee dataset. We computed geographic versus song distance matrices for each ecoregion and song feature pair separately and applied a Mantel test to assess for correlation between geographic distance and song feature distance. An asterisk (*) indicates *p*_adj_ < 0.05 after an FDR correction for multiple comparisons.

However, when we assessed if there was closer association between song and geographic distance within ecoregions compared to the species distribution as a whole in the modern towhee dataset, we found that only average syllable upper frequency in the Great Plains was statistically significant in the Spotted towhees relative to species distribution, and only overall syllable frequency range in the northern forests was statistically significant in the Eastern towhees. In the full towhee dataset, none of the ecoregions for either the Spotted or Eastern towhee were significantly different from the species distribution.

Finally, we tested whether there was a difference in song features between ecoregions when accounting for geographic distance by comparing the distribution of residuals of the distance matrices within each ecoregion to the distribution of residuals of the distance matrices of the entire dataset across all ecoregions for each species. For the Spotted towhees in the modern towhee dataset, we found that 92 out of 168 comparisons were significantly different from the within-region distribution (*p*_adj_ < 0.05; **Fig. S20**), and 16 out of 24 observations in the Eastern towhee were statistically different (*p*_adj_ < 0.05; **Fig. S21**). For example, in the Spotted towhees, we found that largest syllable frequency range in the Southern Semiarid Highlands versus Mediterranean California were more different than expected (mean_residual_ = 0.12; *p*_adj_ = 3.81×10^−28^) relative to the mean residual dissimilarity of within ecoregion pairs. Minimum syllable frequency (mean_residual_ = 0.04; *p*_adj_ = 5.53×10^−57^) in the Eastern towhees between Eastern Temperate Forests and Northern Forests was more different than expected after accounting for geographic distance relative to the dissimilarity between pairs within the same ecoregion. In the full towhee dataset, 190 out of the 224 comparisons for the Spotted towhees were significantly different from the within-region distribution (*p*_adj_ < 0.05; **Fig. S22**), and 42 of 48 comparisons in the Eastern towhee were statistically different (*p*_adj_ < 0.05; **Fig. S23**).

## DISCUSSION

This study provides insight into how ecology, geography, and urbanization can influence the variation in song frequency measures of birdsong in Eastern and Spotted towhees. Overall, the GLM models of both species’ songs depict that species classification has the strongest association with song frequency variables: Eastern and Spotted towhee songs have distinct but overlapping frequency distributions. Further, longitude had considerable additional predictive power for most frequency metrics, likely due to within-species differences across their respective ranges (Spotted towhees in the west and Eastern towhees further east). Our results suggest that songs of Eastern towhees further east have higher frequencies and greater bandwidth. However, the songs of Spotted towhees further east have lower average syllable upper frequency, minimum syllable frequency, smallest syllable frequency range, and average syllable frequency range and higher overall syllable frequency range. These trends, along with the results of the other ecological variables (latitude, urbanization, and elevation) in the GLM, suggest that environmental factors are likely contributing, to some degree, to the variation we see within each species due to certain selection pressures and environmental constraints. For example, in both species, elevation was positively correlated with average syllable upper frequency, maximum syllable frequency, overall syllable frequency range, largest syllable frequency range, and average syllable frequency range. Although previous studies have shown that birds at higher elevations tend to sing at a lower pitch (e.g. green hylia; (Kirschel et al. 2009), it is possible that our observations reflect an increase in pitch as a result of other factors not studied here, such as body size.

Latitude had similar trends in both Spotted and Eastern towhees: going from south to north, both species appear to reduce their upper song frequencies. Additionally, the Eastern towhees further north have lower song frequency in other song traits (maximum syllable frequency, minimum syllable frequency) and narrower bandwidth (largest syllable frequency range, smallest syllable frequency range, and average syllable frequency range). If populations further north have larger body size in accordance with Bergmann’s Rule, (i.e., that individuals in colder climates tend to have a larger body size than individuals in warmer climates closer to the equator) this could contribute to the evolution of lower pitched songs and narrower bandwidths due to morphological constraints (Sagar et al. 2024). Nonetheless, latitude likely reflects underlying ecological patterns that influence frequency variables of song differently across regions, potentially indicating that there are other selection pressures involved in the evolution of its song, such as increased anthropogenic noise, selection for smaller body size, cultural boundaries, or different local environmental pressures. These trends in the variation of song features suggest that more local aspects of ecology (e.g. combination of climate and habitat structure) might contribute to the variation in songs we see in this pair of sister species.

Further, region-specific factors may also contribute to species-specific song evolution. We tested the acoustic adaptation hypothesis in each species, using tree cover as a proxy of foliage density, and we did not see any of the predicted associations with frequency (smaller frequency bandwidth and lower frequencies in areas with greater tree cover), suggesting that acoustic adaptation to dense habitats is not a primary driver of frequency change in towhee songs. Greater levels of urban noise, in contrast, have been linked to elevated frequencies in several species, as birds increase their pitch to avoid low-frequency anthropogenic sounds. Urbanization was associated with Spotted towhee song frequency in the predicted direction: as urbanization increases, so does average syllable lower frequency (negatively associated with distance to road). This suggests that individuals might increase their pitch to prevent the low-frequency anthropogenic noise from masking their songs (e.g. (Pohl et al. 2012; Bermúdez-Cuamatzin et al. 2011; Derryberry et al. 2020). However, in Eastern towhees, population density had a weak but positive association with overall frequency range, but there was no signal of upper or lower frequencies increasing in more urban areas. Overall, these findings show some species-specific divergence in song frequency trends and some shared environmental pressures in both species.

To further assess if climate and ecology contribute to species-specific song variation, we compared the distribution of song frequency features within specific climate zones or ecoregions to the overall species distribution and found that the Spotted towhees showed more significant differences in the song distributions in specific regions than the Eastern towhees. In the warm summer Mediterranean climate zone, frequency range was more narrow and had lower upper frequencies and higher low frequencies in Spotted towhees relative to the entire species distribution, contrasting with humid continental mild summer, wet all year where maximum frequency was higher and contributing to a larger bandwidth. This is consistent with previous literature that suggests that sounds of higher frequency are more easily attenuated by absorption in areas with higher temperatures and lower humidity (Marler and Slabbekoorn 2004); thus, birdsong in the warm summer Mediterranean climate zone, which has warm, dry summers, is expected to have more narrow frequency bandwidths and lower peak frequency compared to songs in the humid continental mild summer, wet all year climate zone, which has more mild, rainy summers and would select for broader frequency bandwidths and higher peak frequency if degradation by attenuation is decreased. Similarly, in the Eastern towhees, average syllable upper frequency, maximum syllable frequency, and overall syllable frequency range was higher than the overall species distribution in the humid continental mild summer, wet all year climate, suggesting that birds might sing at higher frequencies since the humid, rainy summers of this climate zone minimizes sound attenuation or to prevent their songs from being masked by ambient noise, like rain. It is also possible that these climate properties of these regions are conflated with greater anthropogenic noise or other factors.

When we assessed whether there is evidence of isolation-by-distance in songs across the entire distribution and within specific ecoregions, we found that geographic distance does not fully explain the song differences we see in the Spotted and Eastern towhees. However, certain ecoregions showed measurable song differences. For example, Spotted towhees show that maximum syllable frequency and overall syllable frequency range is higher in the Great Plains relative to the overall distribution. Interestingly, for the eastern towhees, song frequency was lower and bandwidth was more narrow in the Great Plains relative to the overall distribution.

The Great Plains have fewer obstacles in the environment due to little topographic relief and a scarcity of forests (**Table S2**; (Commission for Environmental Cooperation, n.d.), providing more room for flexibility in terms of song traits that individuals possess. In other words, song traits, like frequency, are not as limited to a particular spectral space since there aren’t as many obstacles that would degrade the songs. Nonetheless, towhees are likely to be present at lower densities in the Great Plains, which would require individuals to have songs that maximize the distance that the sound can travel to reach another individual. Thus, songs that have a lower pitch might be favored in this environment, as they would travel longer distances and minimize degradation by elements of the environment, particularly wind, which is extremely prevalent in this region.

While our study provides a detailed look into interactions between ecology and song evolution, it does not directly address this interaction at a population level and is limited by the number of variables being examined. Our finding that longitude was the largest contributor to the variation in song frequency measures in the Spotted and Eastern towhees could correspond to non-environmental factors, such as species range and cultural differences along this gradient, that could drive song variation. Other ecological variables, and even morphological constraints not tested here, might also play a role in the song variation given that our analyses also picked up on relatively weak signals from broad analyses of ecoregions, climate zones, latitude, and other variables that reflect, but do not directly measure, specific conditions in habitats. Further, our study was limited to song frequency features, but other song features, such as amplitude, rhythmicity, trill rate, and the number of songs produced per minute, could be more strongly correlated with particular ecological variables (Coomes and Derryberry 2021) or shifts in the local soundscape. For example, white-crowned sparrows were found to sing at a higher performance and lower amplitude in response to the reduction in noise pollution due to the COVID-19 shutdown (Derryberry et al. 2020).

Nonetheless, certain environmental factors do appear to influence some frequency aspects of Eastern and Spotted towhee songs. Specific populations of Spotted and Eastern towhees in regions of the United States may adapt differently to specific environmental factors, so both computational and field studies on local population- and individual-level differences, coupled with data on body size and ecology could shed more light on the local environmental constraints affecting birdsong. In sum, our study provided an analysis of the potential effects of the environment on song evolution in the Spotted and Eastern towhees across a large geographic scale. Overall, we found a stronger relationship between towhee song features and longitude, latitude, and elevation than with our tested metrics of urbanization, environment, or climate, suggesting that the geographic patterns observed in towhee songs could be driven by cultural differences in learned song more than by broad-scale adaptations to ecological or anthropogenic factors. However, these geographic patterns did not lead to a simple pattern of isolation by distance, where nearby songs would be more similar than geographically distant songs. In addition, we did not find support for the broad predictions of the acoustic adaptation hypothesis in these species, but we found a signal that urbanization was associated with increased frequencies in the Spotted towhee. Future studies using additional environmental, cultural, morphological variables, and song features would allow us to better understand what contributes to the song variation in this pair of sister species.

## Author Contributions

Ximena León Du’Mottuchi and Nicole Creanza conceived and designed the project; Ximena León Du’Mottuchi and Meklit Mesfin downloaded and parsed song recordings to produce song-feature data; Ximena León Du’Mottuchi ran all analyses and created the figures and tables; Ximena León Du’Mottuchi and Nicole Creanza revised and edited the manuscript. All authors contributed to the drafts and gave final approval for publication.

## Data and materials availability

All analysis code and data are available at https://github.com/CreanzaLab/TowheeEcology, including metadata about each song recording.

