## Supplemental Materials for "Song evolution in light of ecosystem differences: exploring effects of urbanization and ecology on temporal and frequency traits of Spotted and Eastern towhee songs"

**Supplementary Materials for "Song evolution in light of ecosystem differences: exploring effects of urbanization and ecology on songs of Spotted and Eastern towhees"**

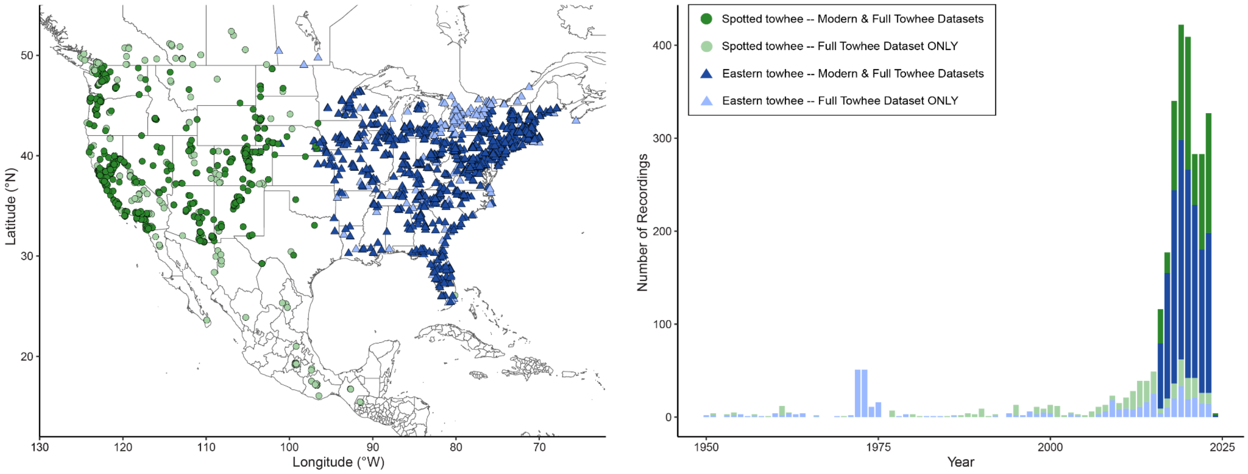


**Supplementary Figure S1.** Geographic and temporal distribution of the Spotted and Eastern towhee song recording samples. **A**) Map showing the geographic distribution of the song recordings across North America. Each point represents an analyzed song in Spotted towhees (*Pipilo maculatus*) and Eastern towhees (*Pipilo erythrophthalmus*). **B**) Barplot showing the number of towhee samples in our analysis grouped by year of recording for the full towhee dataset ($N_{Eastern\_full}$ = 1830; $N_{Spotted\_full}$ = 1086) and the modern towhee dataset ($N_{Eastern\_modern}$ = 1387; $N_{Spotted\_modern}$ = 711). Light blue and light green indicate recordings that were used only in the analyses of the full towhee dataset. Dark blue and dark green indicate the top ~72% of recordings within the continental United States between 2016-2024 that were subsetted from the full dataset and used in both the analyses of the full towhee dataset and the modern towhee dataset.

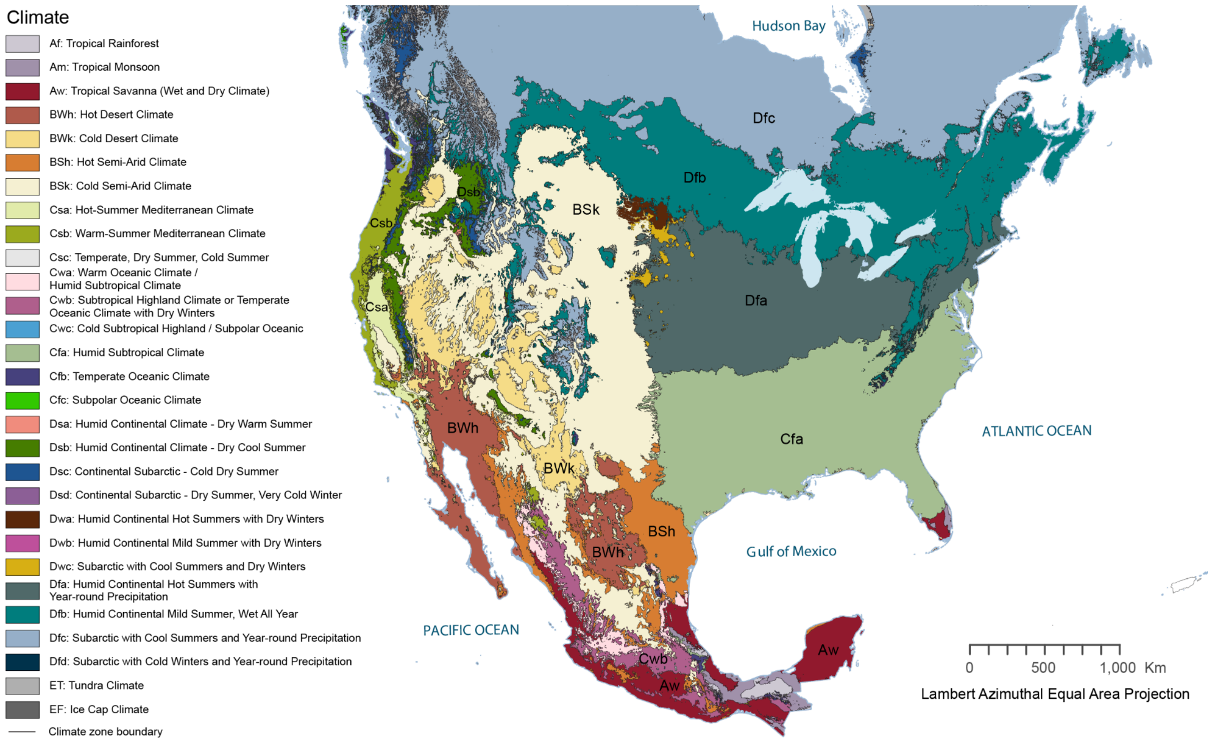


**Supplementary Figure S2.** Map showing Köppen-Geiger climate zones of North America. Adapted from the Commission for Environmental Cooperation (Peel *et al.*, 2007).

| Climate type level 1st | 2nd | 3rd | Description | Criteria | Climate zone label (for zones included in analysis) |
|---|---|---|---|---|---|
| B | | | **Arid** | MAP < 10 x $P_{threshold}$ | — |
| | W | | Desert | MAP < 5 x $P_{threshold}$ | — |
| | | k | Cold | MAT < 18 | BWk: Cold Desert |
| | S | | Steppe | MAP ≥ 5 x $P_{threshold}$ | — |
| | | h | Hot | MAT ≥ 18 | BSh: Hot Semi-Arid |
| | | k | Cold | MAT < 18 | BSk: Cold Semi-Arid |
| C | | | **Temperate** | $T_{hot}$ > 10 & 0 < $T_{cold}$ < 18 | — |
| | s | | Dry Summer | $P_{sdry}$ < 40 & $P_{sdry}$ < $P_{wwet}$ / 3 | — |
| | | a | Hot Summer | $T_{hot}$ ≥ 22 | Csa: Hot Summer Mediterranean |
| | | b | Warm Summer | Not (a) & $T_{mon10}$ ≥ 4 | Csb: Warm Summer Mediterranean |
| | w | | Dry Winter | $P_{wdry}$ < $P_{swet}$ /10 | — |
| | | b | Warm Summer | Not (a) & $T_{mon10}$ ≥ 4 | Cwb: Subtropical Highland/ Temperate Oceanic with Dry Winters |
| | f | | Without dry season | Not (Cs) or (Cw) | — |
| | | a | Hot Summer | $T_{hot}$ ≥ 22 | Cfa: Humid Subtropical |
| | | b | Warm Summer | Not (a) & $T_{mon10}$ ≥ 4 | Cfb: Temperate Oceanic |
| D | | | **Cold** | $T_{hot}$ > 10 & $T_{cold}$ ≤ 0 | — |
| | s | | Dry Summer | $P_{sdry}$ < 40 & $P_{sdry}$ < $P_{wwet}$ / 3 | — |
| | | b | Warm Summer | Not (a) & $T_{mon10}$ > 4 | Dsb: Humid Continental - Dry Cool Summer |
| | f | | Without dry season | Not (Ds) or (Dw) | — |
| | | a | Hot Summer | $T_{hot}$ ≥ 22 | Dfa: Humid Continental Hot Summers with Year-round Precipitation |
| | | b | Warm Summer | Not (a) & $T_{mon10}$ > 4 | Dfb: Humid Continental Mild Summer, Wet All Year |

**Supplementary Table S1.** Descriptions of the Köppen-Geiger climate zones of North America, with criteria and abbreviations adapted from (Peel et al., 2007): MAP = mean annual precipitation, MAT = mean annual temperature, $T_{hot}$ = temperature of the hottest month, $T_{cold}$ = temperature of the coldest month, $T_{mon10}$ = number of months where the temperature is above 10, $P_{dry}$ = precipitation of the driest month, $P_{sdry}$ = precipitation of the driest month in summer, $P_{wdry}$ = precipitation of the driest month in winter, $P_{swet}$ = precipitation of the wettest month in summer, $P_{wwet}$ = precipitation of the wettest month in winter, $P_{threshold}$ = varies according to the following rules (if 70% of MAP occurs in winter then $P_{threshold}$ = 2 x MAT, if 70% of MAP occurs in summer then $P_{threshold}$ = 2 x MAT + 28, otherwise $P_{threshold}$ = 2 x MAT + 14). Summer (winter) is defined as the warmer (cooler) six month period of ONDJFM and AMJJAS.

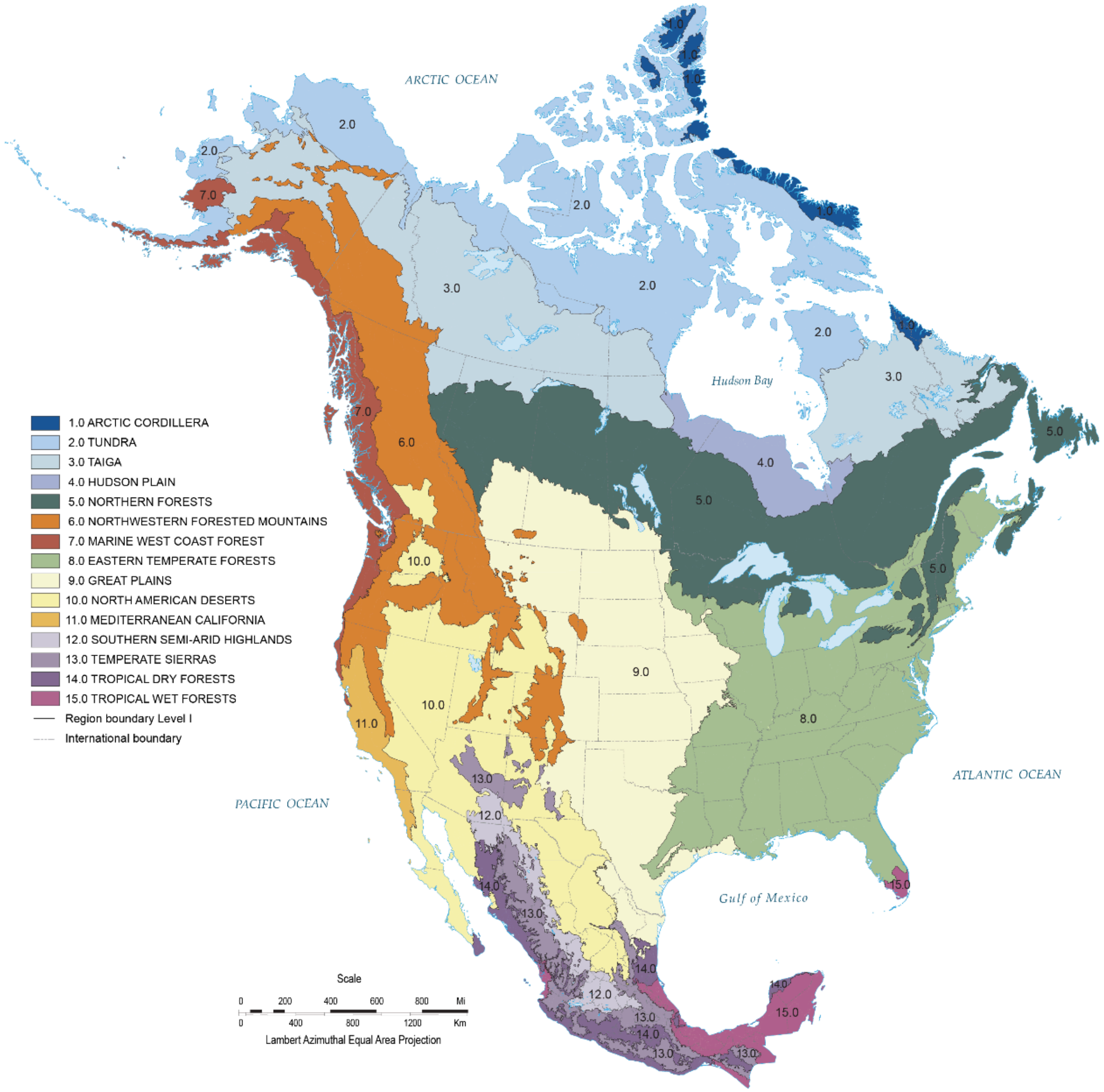


**Supplementary Figure S3.** Map showing Level I Ecoregions of North America. Adapted from the Commission for Environmental Cooperation (*Ecological Regions of North America: Toward a Common Perspective*, 1997).

**Supplementary Table S2.** Summary of Level I Ecoregions of North America.

| Ecoregion | Description |
|---|---|
| Arctic Cordillera | Rugged mountain chain with extensive icefields and glaciers |
| Tundra | Cold, treeless plains with permafrost and short growing seasons |
| Taiga | Boreal forest with conifer-dominated cover and long winters |
| Hudson Plain | Flat, wetland-rich lowlands around Hudson Bay |
| Northern Forests | Mixed needleleaf and broadleaf forests across central Canada |
| Northwestern Forested Mountains | Mountainous terrain with varied forests and alpine areas |
| Marine West Coast Forests | Moist temperate forests along the Pacific coast with high precipitation |
| Eastern Temperate Forests | Broadleaf deciduous forests and mixed woodlands in Eastern North America |
| Great Plains | Flat to rolling grasslands and prairies with continental climate |
| North American Deserts | Plains with hills, plains with mountains, and tablelands of high relief with arid to semi-arid climates with seasonal temperature extremes |
| Mediterranean California | Coastal and inland areas with hot, dry summers and cool, wet winters—dominated by chaparral and oak woodlands |
| Southern Semi-Arid Highlands | Mountainous terrain in Mexico with oak-pine forests and greater moisture than nearby deserts. |
| Temperate Sierras | Elevationally diverse mountains with temperate and subtropical montane vegetation |
| Tropical Dry Forests | Seasonal dry forests in Mexico & Yucatán—deciduous trees adapted to long dry seasons. |
| Tropical Wet Forests | Humid, evergreen and semideciduous forests in southern and western Mexico, Yucatán, and southern Florida. |

Brief descriptions of the fifteen continental-scale ecological regions defined by the Commission for Environmental Cooperation (*Ecological Regions of North America: Toward a Common Perspective*, 1997), characterized by broad differences in climate, vegetation, and landforms.

**Table S3.** Generalized linear model of ecological variables versus song frequency variables for Spotted and Eastern towhees separately in full towhee dataset.

| | | Longitude | | Latitude | | Population Density | | Nighttime Lights | | Distance to Road | | Tree Cover | | Elevation | |
|---|---|---|---|---|---|---|---|---|---|---|---|---|---|---|---|
| | | z | $p_{adj}$ | z | $p_{adj}$ | z | $p_{adj}$ | z | $p_{adj}$ | z | $p_{adj}$ | z | $p_{adj}$ | z | $p_{adj}$ |
| Average Syllable Upper Frequency | S | -2.57 | **0.01** | -4.02 | **$6.22x10^{-5}$** | -0.15 | 0.88 | 1.93 | 0.05 | -2.05 | **0.04** | -0.86 | 0.39 | 3.32 | **$9.46x10^{-4}$** |
| | E | 7.39 | **$2.32x10^{-13}$** | -3.37 | **$7.73x10^{-4}$** | 1.89 | 0.06 | -1.50 | 0.13 | 0.84 | 0.40 | 1.07 | 0.28 | 2.27 | **0.02** |
| Average Syllable Lower Frequency | S | -1.06 | 0.29 | -0.70 | 0.48 | -1.42 | 0.15 | 3.47 | **$5.42x10^{-4}$** | -1.34 | 0.18 | -0.89 | 0.37 | 1.70 | 0.09 |
| | E | 4.26 | **$2.16x10^{-5}$** | 0.30 | 0.77 | 1.83 | 0.07 | -0.51 | 0.61 | 0.43 | 0.67 | 1.86 | 0.06 | 1.28 | 0.20 |
| Maximum Syllable Frequency | S | 4.37 | **$1.38x10^{-5}$** | -1.24 | 0.22 | -0.91 | 0.36 | 2.13 | **0.03** | -1.74 | 0.08 | -0.07 | 0.95 | 6.10 | **$1.45x10^{-9}$** |
| | E | 6.28 | **$4.27x10^{-10}$** | -3.95 | **$8.17x10^{-5}$** | 1.65 | 0.10 | -2.68 | **$7.38x10^{-3}$** | -0.15 | 0.88 | 0.07 | 0.94 | 3.04 | **$2.43x10^{-3}$** |
| Minimum Syllable Frequency | S | -4.95 | **$8.51x10^{-7}$** | 2.22 | **0.03** | -1.36 | 0.17 | 0.82 | 0.41 | -0.22 | 0.82 | -0.41 | 0.68 | -2.26 | 0.02 |
| | E | 0.84 | 0.40 | -4.81 | **$1.66x10^{-6}$** | 1.58 | 0.11 | -1.04 | 0.30 | -0.63 | 0.53 | -0.35 | 0.73 | -0.93 | 0.35 |
| Overall Syllable Frequency Range | S | 6.52 | **$1.10x10^{-10}$** | -1.91 | 0.06 | -0.75 | 0.45 | 1.68 | 0.09 | -1.25 | 0.21 | -0.13 | 0.90 | 7.20 | **$1.11x10^{-12}$** |
| | E | 5.67 | **$1.68x10^{-8}$** | -2.05 | **0.04** | 1.01 | 0.31 | -2.19 | **0.03** | 0.07 | 0.95 | 0.22 | 0.83 | 3.30 | **$9.86x10^{-4}$** |
| Largest Syllable Frequency Range | S | 2.17 | **0.03** | -2.72 | **$6.70x10^{-3}$** | 1.06 | 0.29 | -0.84 | 0.40 | -0.74 | 0.46 | 0.31 | 0.76 | 1.91 | 0.06 |
| | E | 3.78 | **$1.60x10^{-4}$** | -3.97 | **$7.32x10^{-5}$** | 0.43 | 0.67 | -1.24 | 0.21 | -0.64 | 0.52 | -0.72 | 0.47 | 0.89 | 0.37 |
| Smallest Syllable Frequency Range | S | -4.16 | **$3.37x10^{-5}$** | -2.07 | **0.04** | 0.74 | 0.46 | -0.54 | 0.59 | -0.60 | 0.55 | 0.95 | 0.34 | 0.85 | 0.39 |
| | E | 3.88 | **$1.08x10^{-4}$** | -3.70 | **$2.25x10^{-4}$** | -0.89 | 0.37 | 0.08 | 0.94 | 1.61 | 0.11 | -0.93 | 0.35 | 0.18 | 0.86 |
| Average Syllable Frequency Range | S | -1.47 | 0.14 | -2.99 | **$2.85x10^{-3}$** | 1.30 | 0.19 | -1.10 | 0.27 | -0.80 | 0.42 | -0.03 | 0.98 | 1.68 | 0.09 |
| | E | 4.29 | **$1.91x10^{-5}$** | -3.68 | **$2.37x10^{-4}$** | 0.61 | 0.55 | -1.14 | 0.25 | 0.52 | 0.60 | -0.21 | 0.83 | 1.29 | 0.20 |

A generalized linear model was applied to raw song feature data of the Eastern towhee (*Pipilo erythrophthalmus*) [$N_{Eastern}$ = 1781] and the Spotted towhee (*Pipilo maculatus*) [$N_{Spotted}$ = 1067] using a Gaussian family log link function with scaled ecological variables. Adjusted *p*-values are shown; results with $p_{adj} < 0.05$ are bolded and highlighted in red.

**Supplementary Table S4.** Generalized linear model of ecological variables versus song frequency variables in full towhee dataset.

| | Species | | Longitude | | Latitude | | Population Density | | Nighttime Lights | | Distance to Road | | Tree Cover | | Elevation | |
|---|---|---|---|---|---|---|---|---|---|---|---|---|---|---|---|---|
| | t | $p_{adj}$ | t | $p_{adj}$ | t | $p_{adj}$ | t | $p_{adj}$ | t | $p_{adj}$ | t | $p_{adj}$ | t | $p_{adj}$ | t | $p_{adj}$ |
| Average Syllable Upper Frequency | 12.89 | **$5.16x10^{-37}$** | 3.64 | **$2.80x10^{-4}$** | -3.42 | **$6.38x10^{-4}$** | 1.11 | 0.27 | -0.01 | 0.99 | -1.68 | 0.09 | 1.33 | 0.18 | 0.77 | 0.44 |
| Average Syllable Lower Frequency | 5.30 | **$1.22x10^{-7}$** | 2.31 | **0.02** | 0.68 | 0.50 | -0.10 | 0.92 | 2.11 | 0.04 | -1.50 | 0.13 | 1.19 | 0.23 | 0.63 | 0.53 |
| Maximum Syllable Frequency | 8.10 | **$7.86x10^{-16}$** | 7.18 | **$8.57x10^{-13}$** | -3.10 | **$1.95x10^{-3}$** | 0.54 | 0.59 | -1.37 | 0.17 | -2.10 | **0.04** | -0.14 | 0.89 | 5.89 | **$4.27x10^{-9}$** |
| Minimum Syllable Frequency | 5.67 | **$1.60x10^{-8}$** | -4.79 | **$1.75x10^{-6}$** | 0.43 | 0.67 | -0.89 | 0.37 | 0.41 | 0.68 | -0.57 | 0.57 | -0.07 | 0.94 | -6.81 | **$1.20x10^{-11}$** |
| Overall Syllable Frequency Range | 5.21 | **$2.03x10^{-7}$** | 8.98 | **$4.78x10^{-19}$** | -3.06 | **$2.23x10^{-3}$** | 0.46 | 0.64 | -1.39 | 0.16 | -1.61 | 0.11 | -0.17 | 0.86 | 8.68 | **$6.61x10^{-18}$** |
| Largest Syllable Frequency Range | 6.65 | **$3.54x10^{-11}$** | 4.13 | **$3.78x10^{-5}$** | -4.49 | **$7.39x10^{-6}$** | 0.98 | 0.33 | -1.52 | 0.13 | -0.93 | 0.35 | -0.42 | 0.68 | 1.45 | 0.15 |
| Smallest Syllable Frequency Range | 7.00 | **$3.08x10^{-12}$** | -3.60 | **$3.27x10^{-4}$** | -2.50 | **0.01** | 0.68 | 0.49 | -0.70 | 0.49 | -0.24 | 0.81 | 1.47 | 0.14 | -1.03 | 0.30 |
| Average Syllable Frequency Range | 7.91 | **$3.71x10^{-15}$** | 1.43 | 0.15 | -3.67 | **$2.52x10^{-4}$** | 1.47 | 0.14 | -1.79 | 0.07 | -0.53 | 0.60 | 0.43 | 0.67 | 0.31 | 0.76 |

A generalized linear model was applied to raw song feature data of the Eastern Towhee (*Pipilo erythrophthalmus*) [$N_{Eastern}$ = 1781] and the Spotted Towhee (*Pipilo maculatus*) [$N_{Spotted}$ = 1067] using a Gaussian family log link function with scaled ecological variables. After multiple hypothesis correction with FDR, results with $p_{adj} < 0.05$ are bolded and highlighted in red.

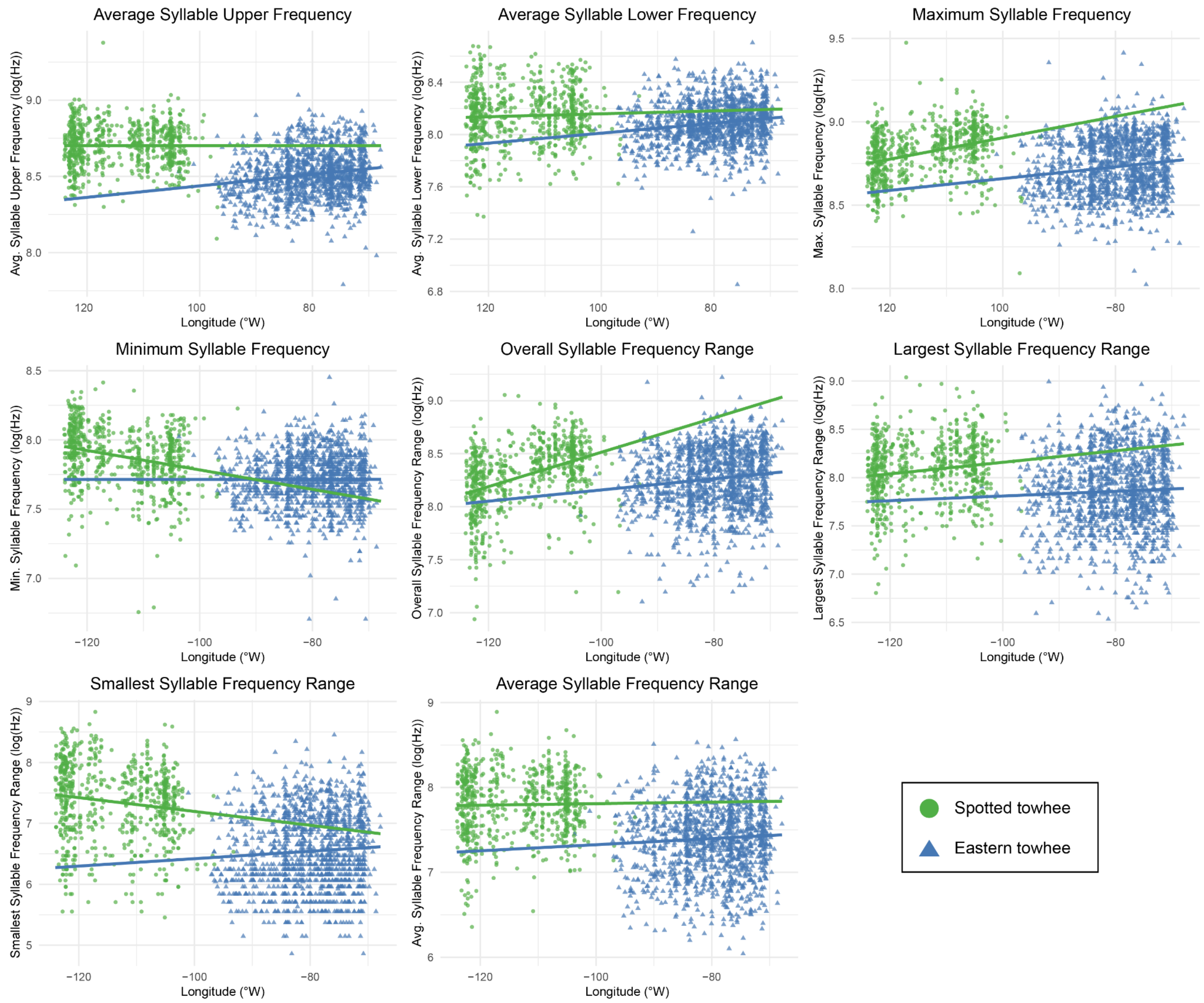


**Supplementary Figure S4.** Scatterplots showing the relationship between log-transformed song frequency features (log(Hz)) and longitude (°W). Each point represents an analyzed song in Eastern towhees (*Pipilo erythrophthalmus*) [N = 1387] and Spotted towhees (*Pipilo maculatus*) [N = 711] from the modern towhee dataset. A linear model was fit to the data from each species separately (indicated by lines in the corresponding color); see **Table 1** for *t* and *p*-values from generalized linear model.

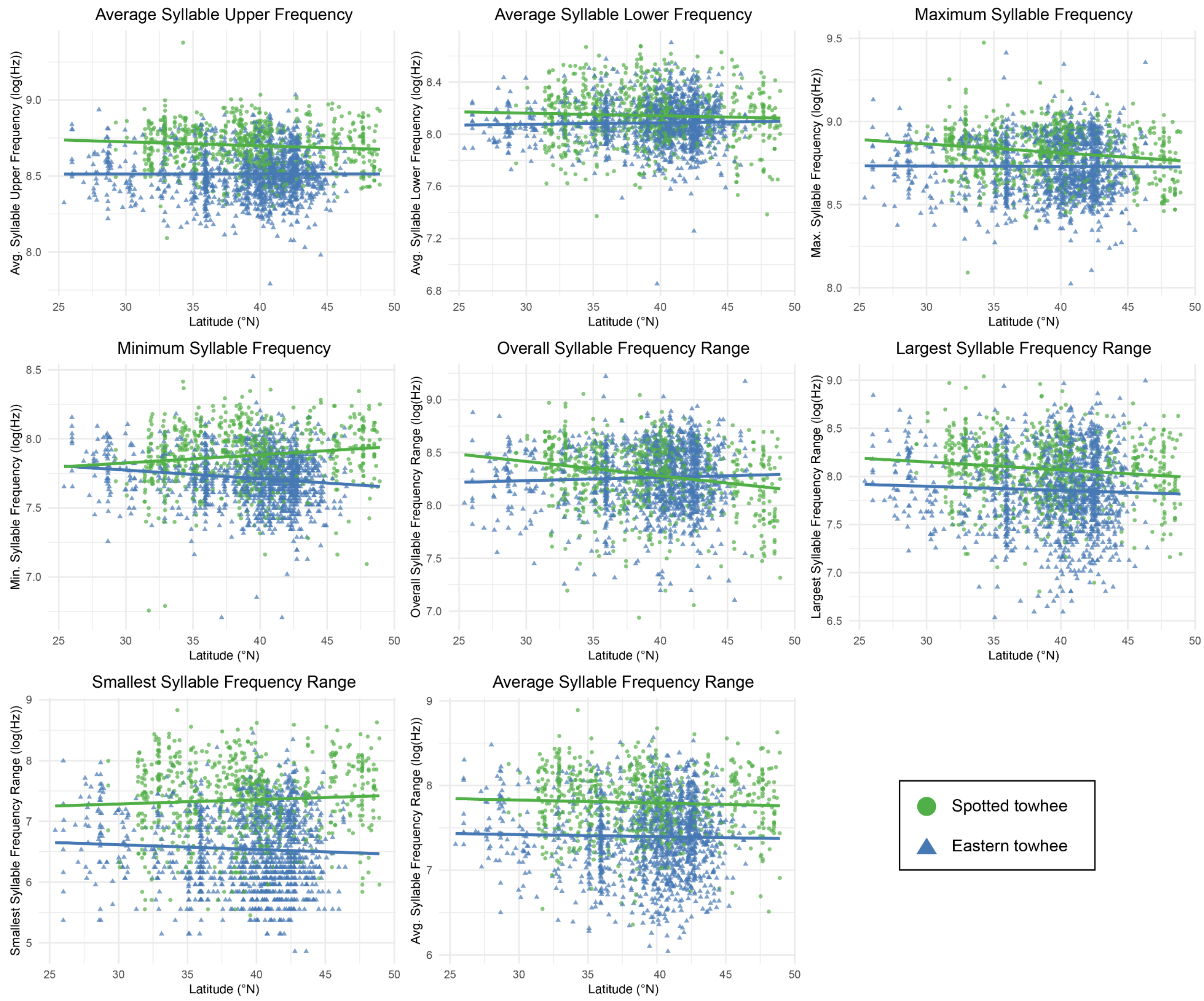


**Supplementary Figure S5.** Scatterplots showing the relationship between log-transformed song frequency features (log(Hz)) and latitude (°N). Each point represents an analyzed song in Eastern Towhees (*Pipilo erythrophthalmus*) [N = 1387] and Spotted Towhees (*Pipilo maculatus*) [N = 711] from the modern towhee dataset. A linear model was fit to the data from each species separately (indicated by lines in the corresponding color); see **Table 1** for *t* and *p*-values from generalized linear model.

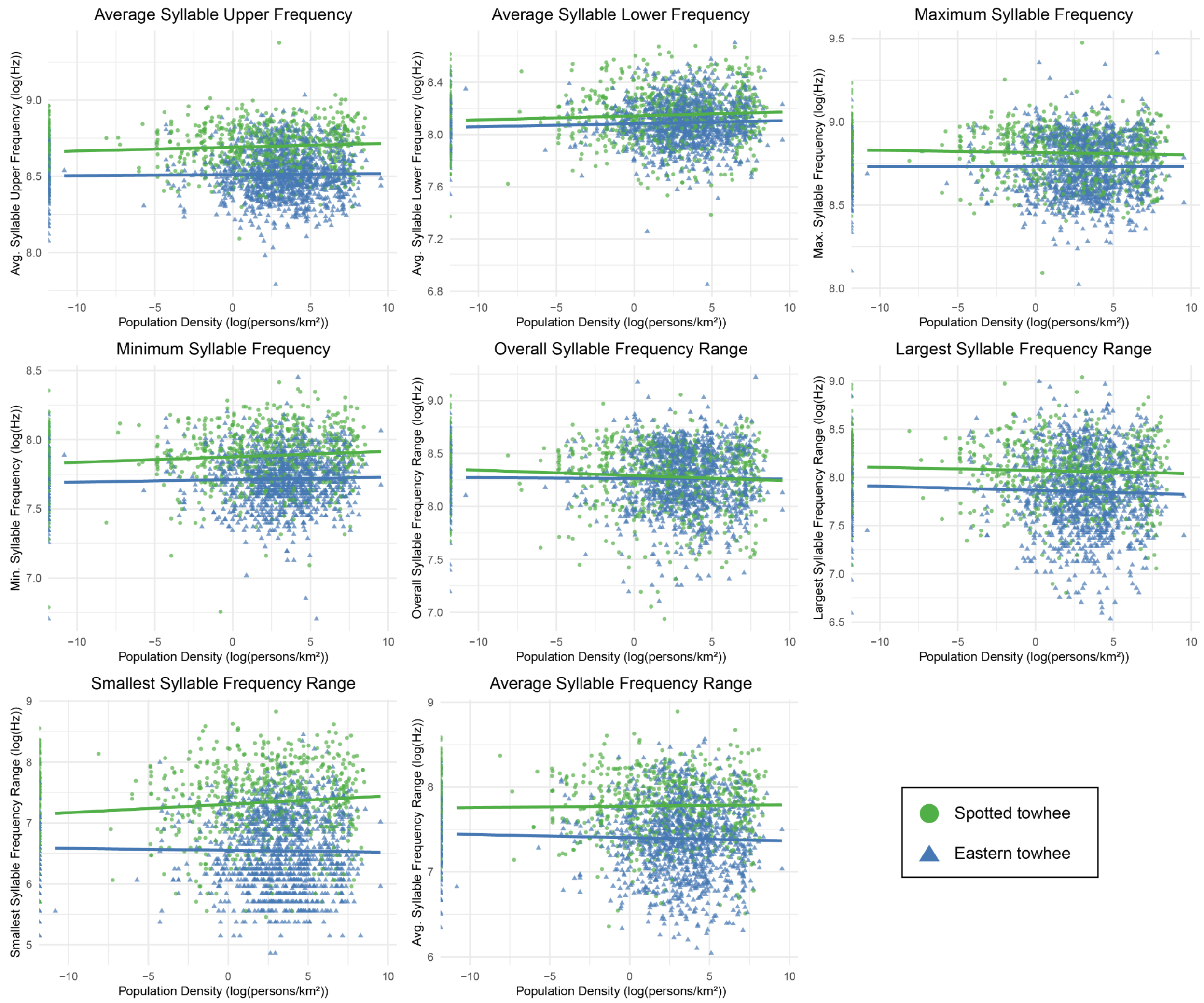


**Supplementary Figure S6.** Scatterplots showing the relationship between log-transformed song frequency features (log(Hz)) and log-transformed population density (log(persons/km$^2$)). Each point represents an analyzed song in Eastern Towhees (*Pipilo erythrophthalmus*) [N = 1387] and Spotted Towhees (*Pipilo maculatus*) [N = 711] from the modern towhee dataset. A linear model was fit to the data from each species separately (indicated by lines in the corresponding color); see **Table 1** for *t* and *p*-values from generalized linear model.

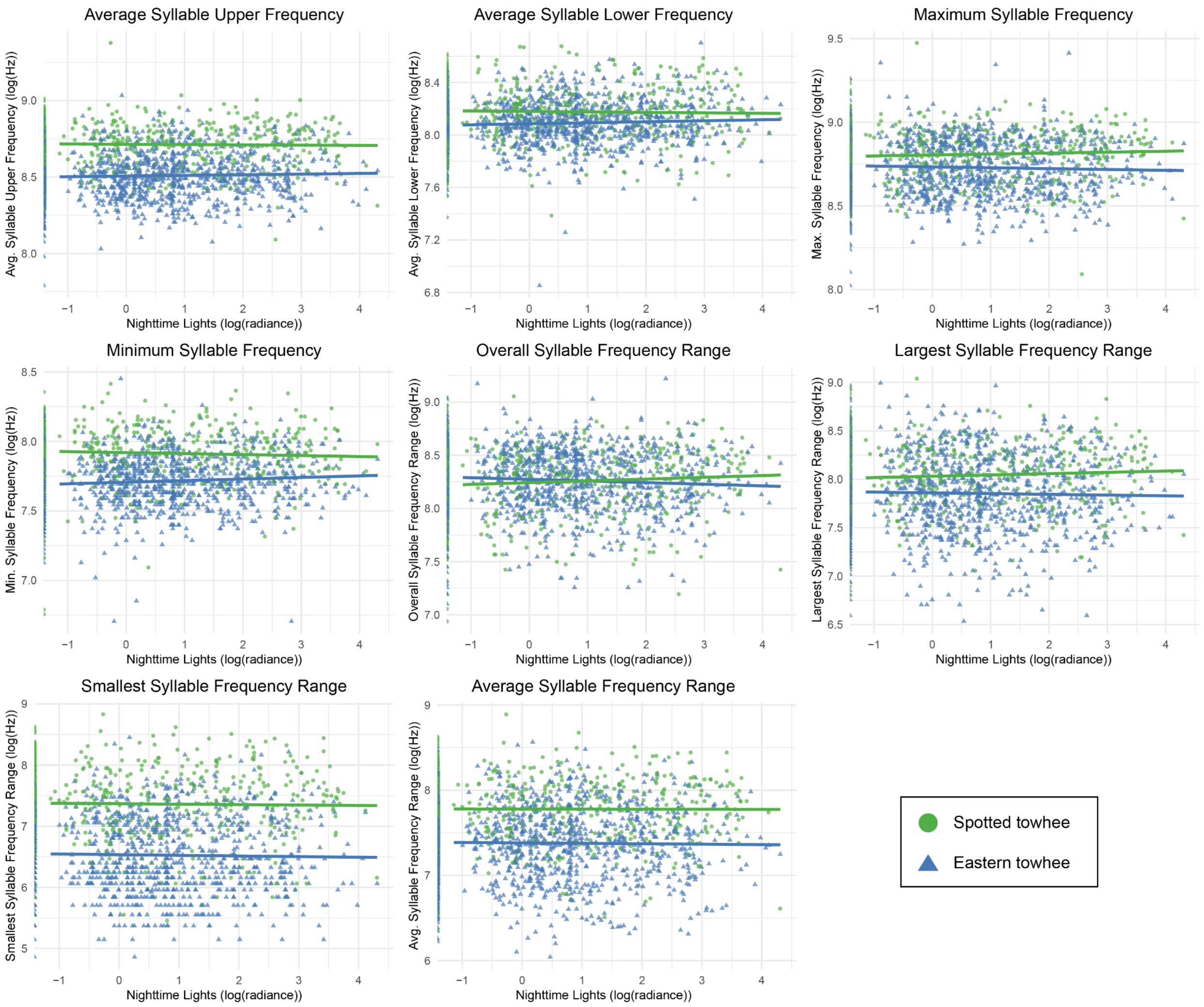


**Supplementary Figure S7.** Scatterplots showing the relationship between log-transformed song frequency features (log(Hz)) and log-transformed stable nighttime light intensity (log(radiance)). Each point represents an analyzed song in Eastern Towhee (*Pipilo erythrophthalmus*) [N = 1387] and Spotted Towhee (*Pipilo maculatus*) [N = 711] from the modern towhee dataset. A linear model was fit to the data from each species separately (indicated by lines in the corresponding color); see **Table 1** for *t* and *p*-values from the generalized linear model.

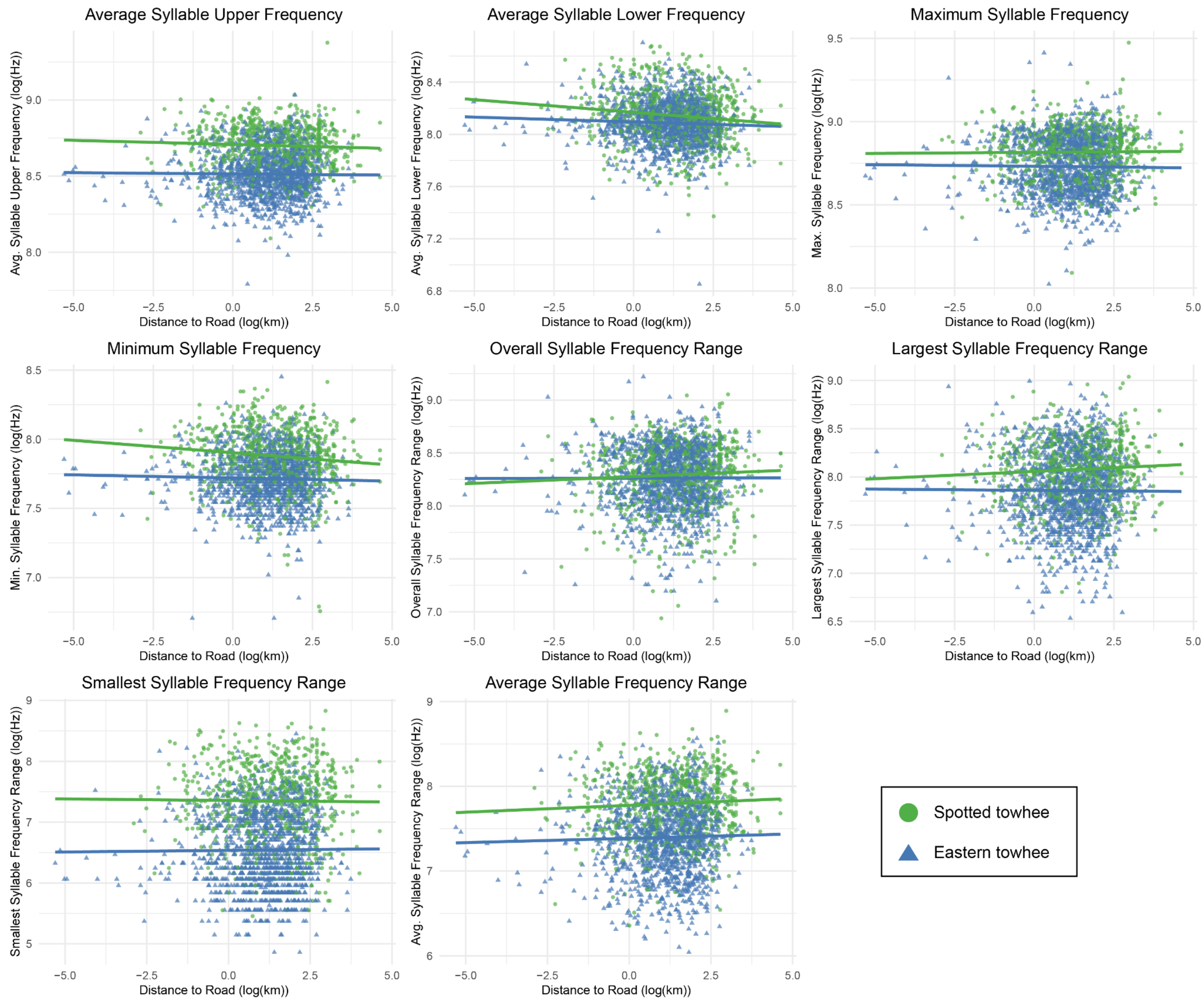


**Supplementary Figure S8.** Scatterplots showing the relationship between log-transformed song frequency features (log(Hz)) and log-transformed distance to road (log(km)). Each point represents an analyzed song in Eastern Towhees (*Pipilo erythrophthalmus*) [N = 1387] and Spotted Towhees (*Pipilo maculatus*) [N = 711] from the modern towhee dataset. A linear model was fit to the data from each species separately (indicated by lines in the corresponding color); see **Table 1** for *t* and *p*-values from the generalized linear model.

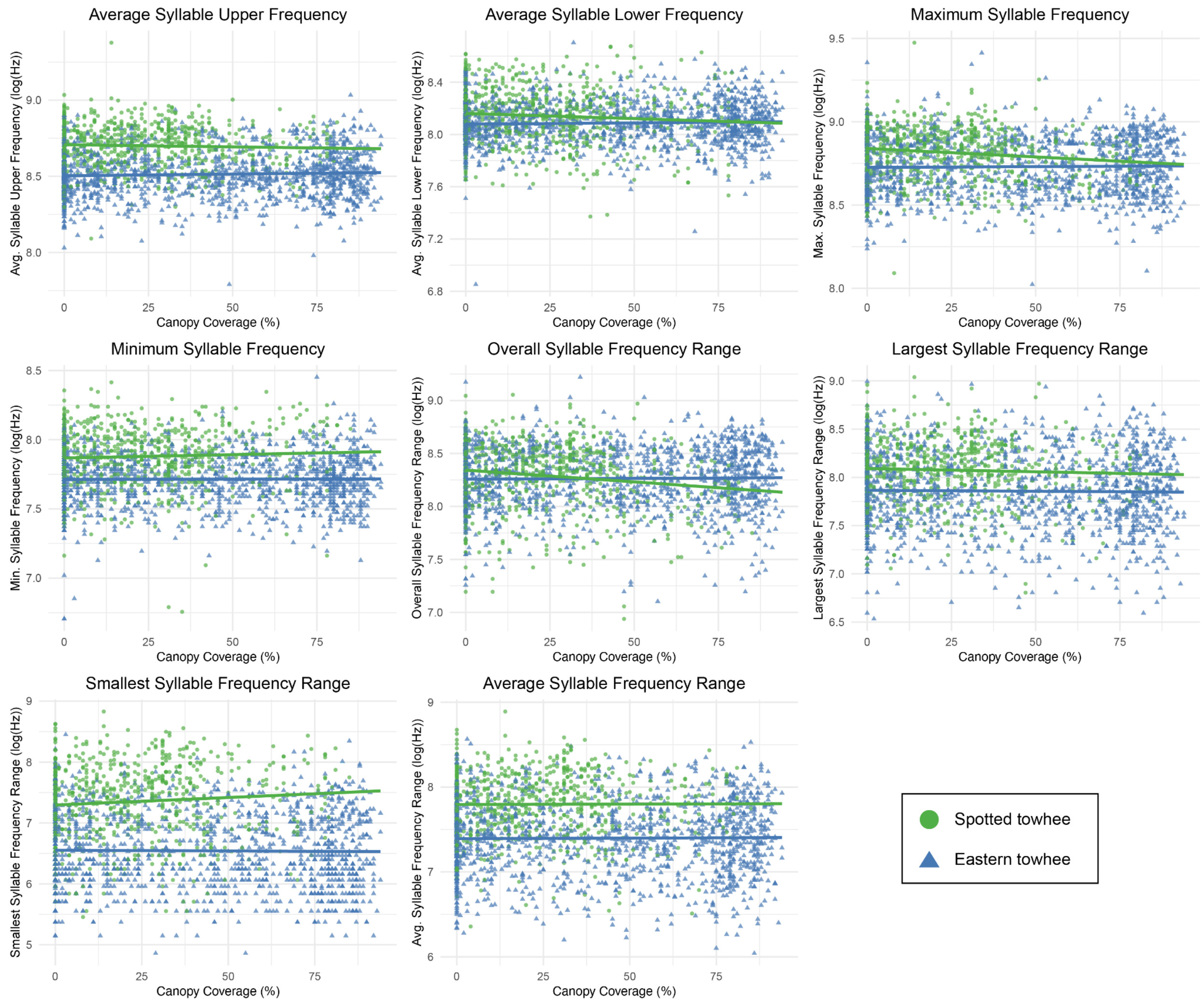


**Supplementary Figure S9.** Scatterplots showing the relationship between log-transformed song frequency features (log(Hz)) and percent canopy coverage. Each point represents an analyzed song in Eastern Towhees (*Pipilo erythrophthalmus*) [N = 1387] and Spotted Towhees (*Pipilo maculatus*) [N = 711] from the modern towhee dataset. A linear model was fit to the data from each species separately (indicated by lines in the corresponding color); see **Table 1** for *t* and *p*-values from the generalized linear model.

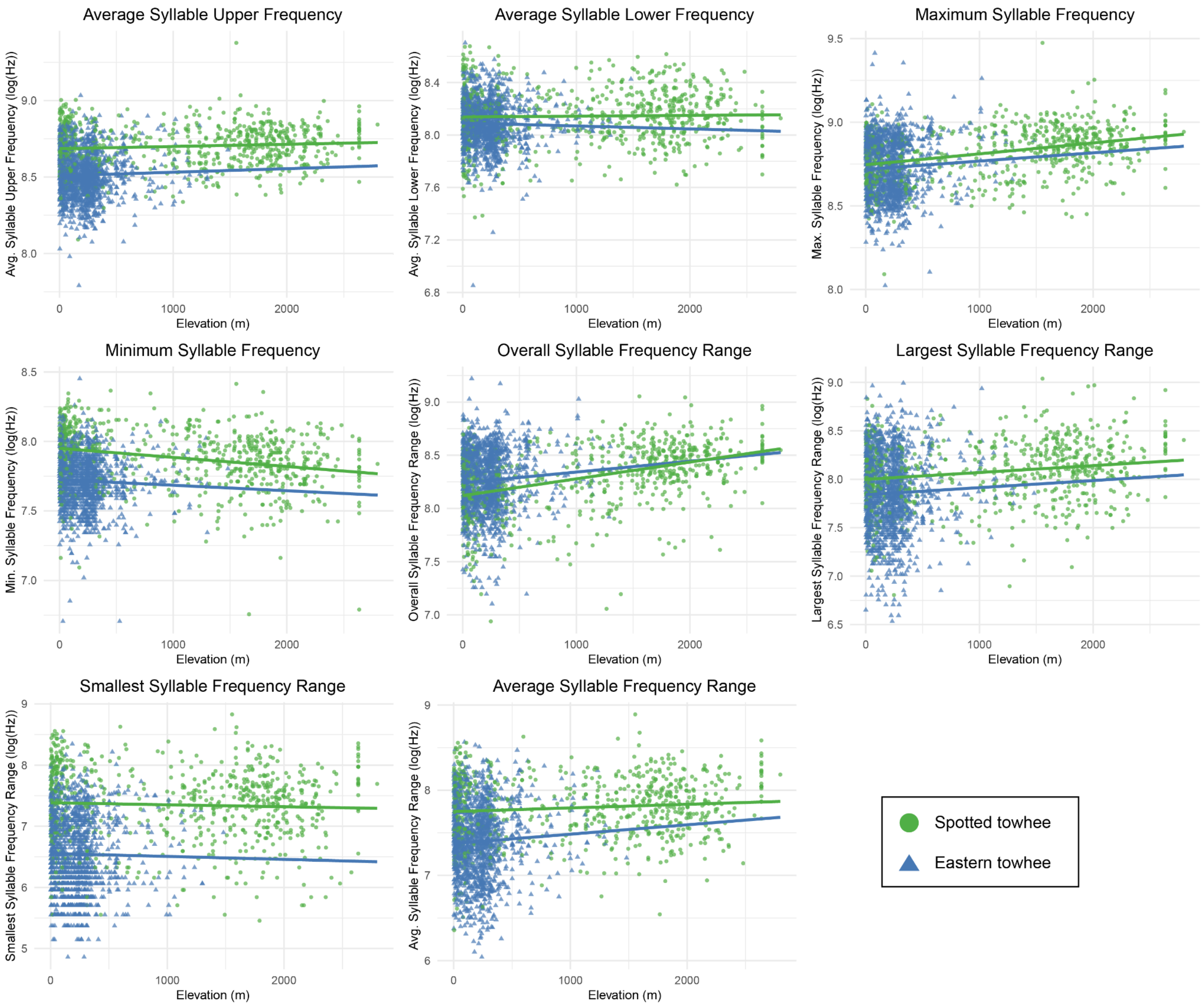


**Supplementary Figure S10.** Scatterplots showing the relationship between log-transformed song frequency features (log(Hz)) and elevation (m). Each point represents an analyzed song in Eastern Towhees (*Pipilo erythrophthalmus*) [N = 1358] and Spotted Towhees (*Pipilo maculatus*) [N = 710] from the modern towhee dataset. A linear model was fit to the data from each species separately (indicated by lines in the corresponding color); see **Table 1** for *t* and *p*-values from the generalized linear model.

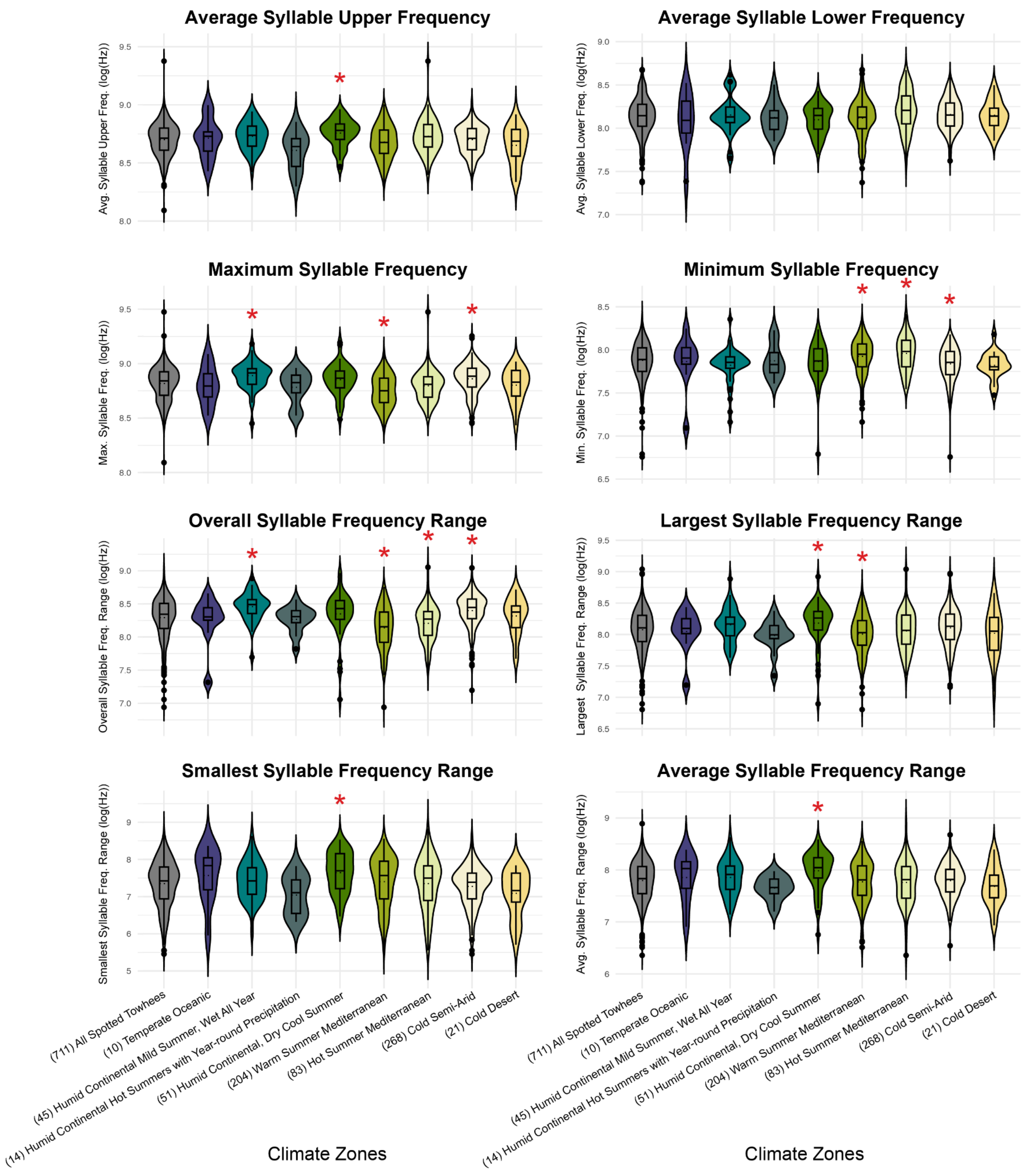


**Supplementary Figure S11.** Violin plots showing the distribution of song frequency traits in each climate zone for the Spotted towhee in the modern towhee dataset. We used Wilcoxon rank-sum tests to assess if there is a statistically significant difference in song frequency traits in each climate zone relative to the overall species distribution (N = 711). The left-most violin on each plot ("All Spotted Towhees") indicates the overall species distribution. An asterisk (*)

indicates $p_{adj} < 0.05$ after adjusting for multiple comparisons using the false discovery rate method.

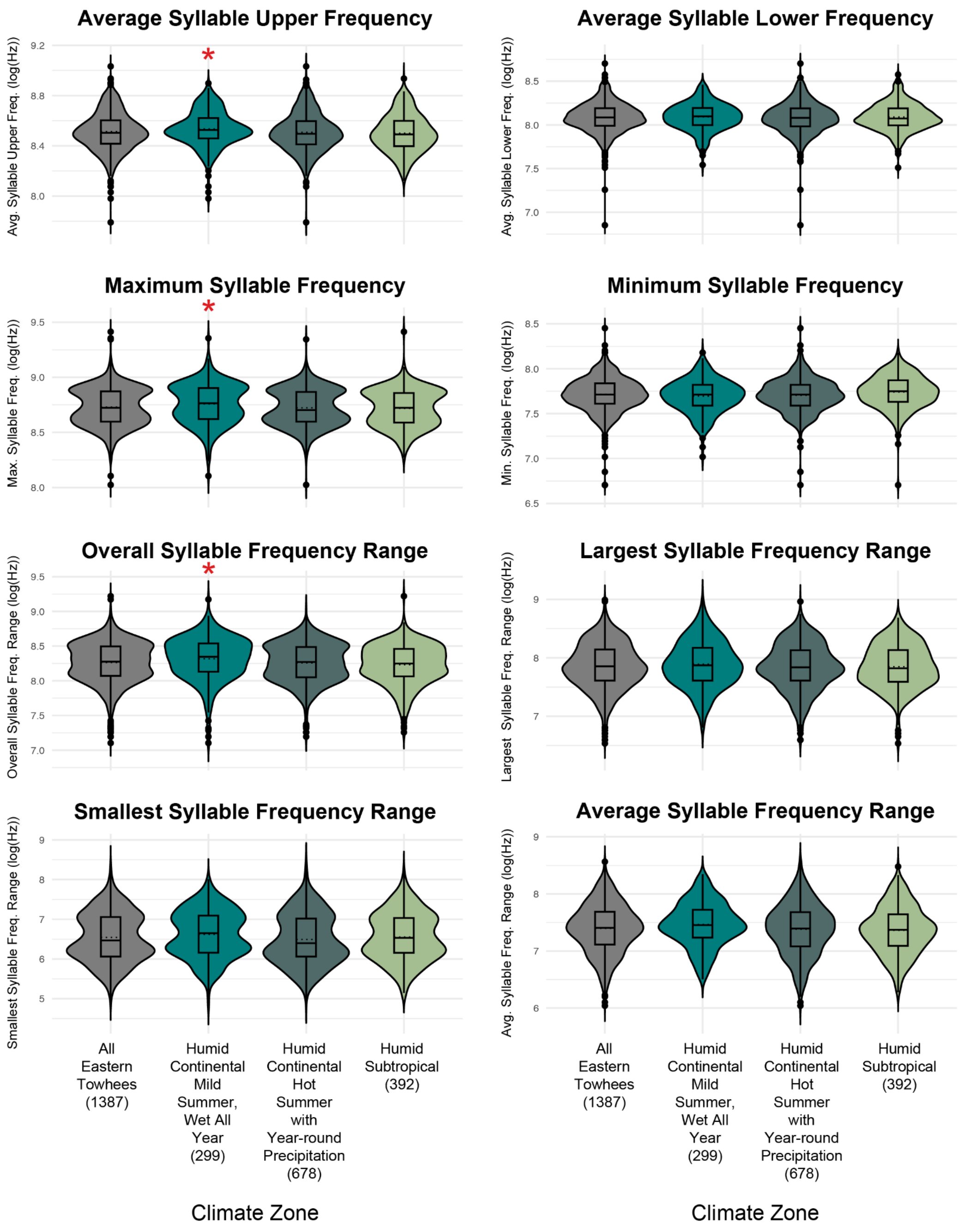


**Supplementary Figure S12.** Violin plots showing the distribution of song frequency traits in each climate zone for the Eastern towhee in the modern towhee dataset. We used a Wilcoxon rank-sum test to assess if there is a statistically significant difference in song frequency traits in each climate zone relative to the overall species distribution. Left-most violin on each plot ("All

Eastern Towhees”) indicates the overall species distribution (N = 1387). An asterisk (*) indicates $p_{adj} < 0.05$ after adjusting for multiple comparisons using the false discovery rate method.

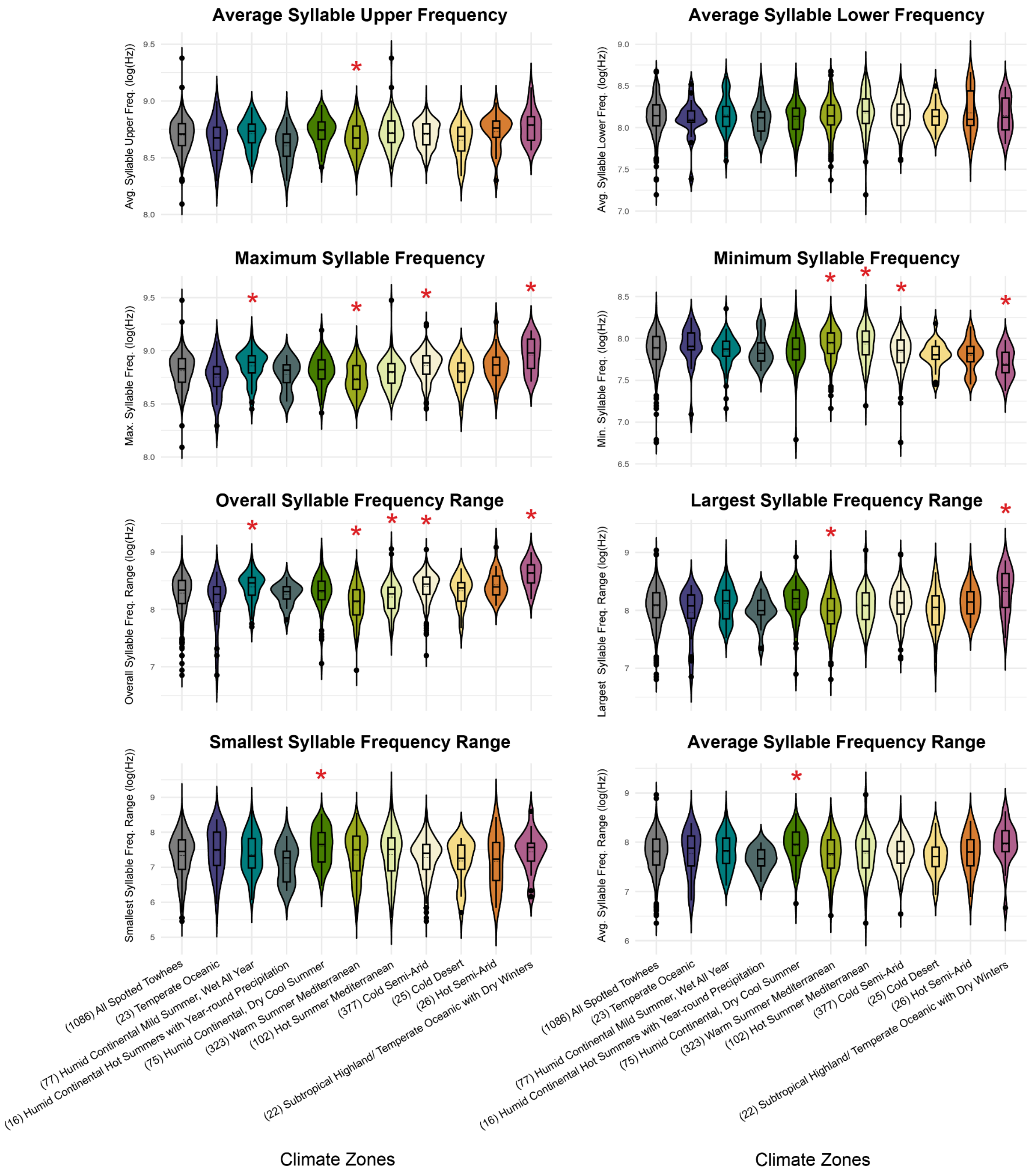


**Supplementary Figure S13.** Violin plots showing the distribution of song frequency traits in each climate zone for the Spotted towhee in the full towhee dataset. We used Wilcoxon rank-sum tests to assess if there is a statistically significant difference in song frequency traits in each climate zone relative to the overall species distribution (N = 1086). The left-most violin on each plot (“All Spotted Towhees”) indicates the overall species distribution. An asterisk (*) indicates $p_{adj} < 0.05$ after adjusting for multiple comparisons using the false discovery rate method.

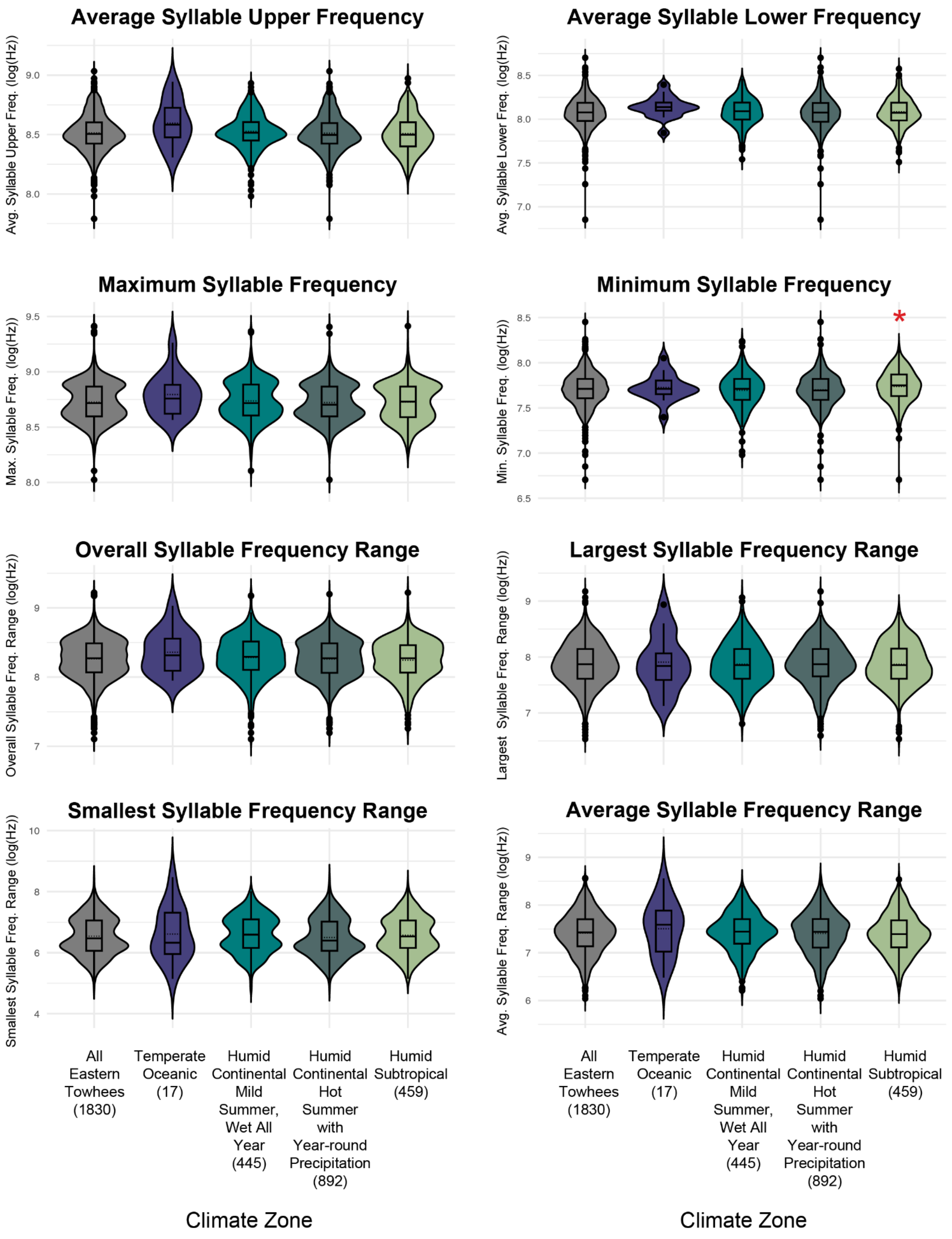


**Supplementary Figure S14.** Violin plots showing the distribution of song frequency traits in each climate zone for the Eastern towhee in the full towhee dataset. We used a Wilcoxon rank-sum test to assess if there is a statistically significant difference in song frequency traits in each climate zone relative to the overall species distribution. Left-most violin on each plot ("All Eastern Towhees") indicates the overall species distribution (N = 1830). An asterisk (*) indicates $p_{adj} < 0.05$ after adjusting for multiple comparisons using the false discovery rate method.

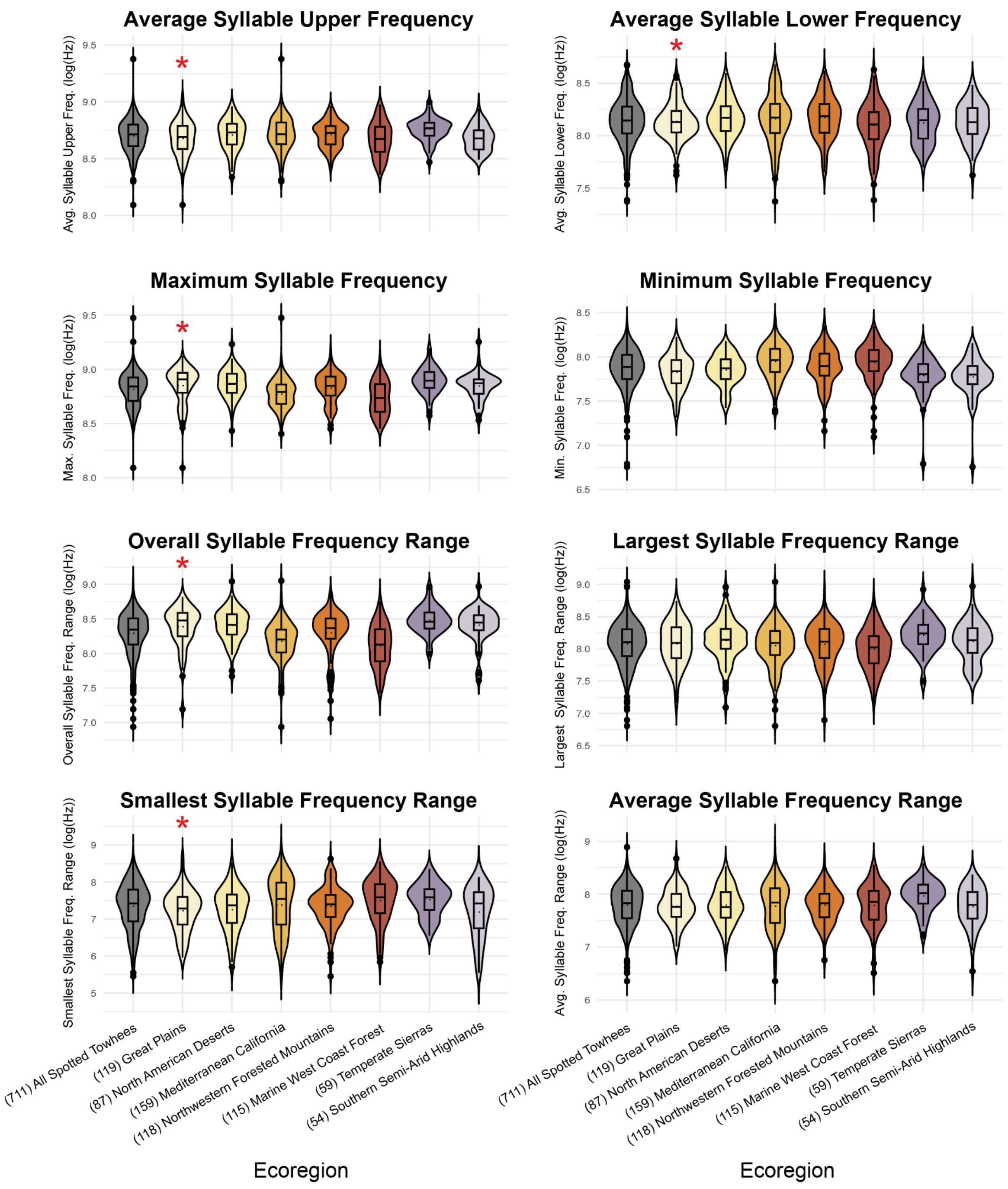


**Supplementary Figure S15.** Violin plots showing the distribution of song frequency traits in each ecoregion for the Spotted towhee in the modern towhee dataset. We used a Wilcoxon rank-sum test to assess if there is a statistically significant difference in song frequency traits in each ecoregion relative to the overall species distribution (N = 711). Left-most violin on each plot (“All Spotted Towhees”) indicates the overall species distribution. An asterisk (*) indicates $p_{adj} < 0.05$ after adjusting for multiple comparisons using the false discovery rate method.

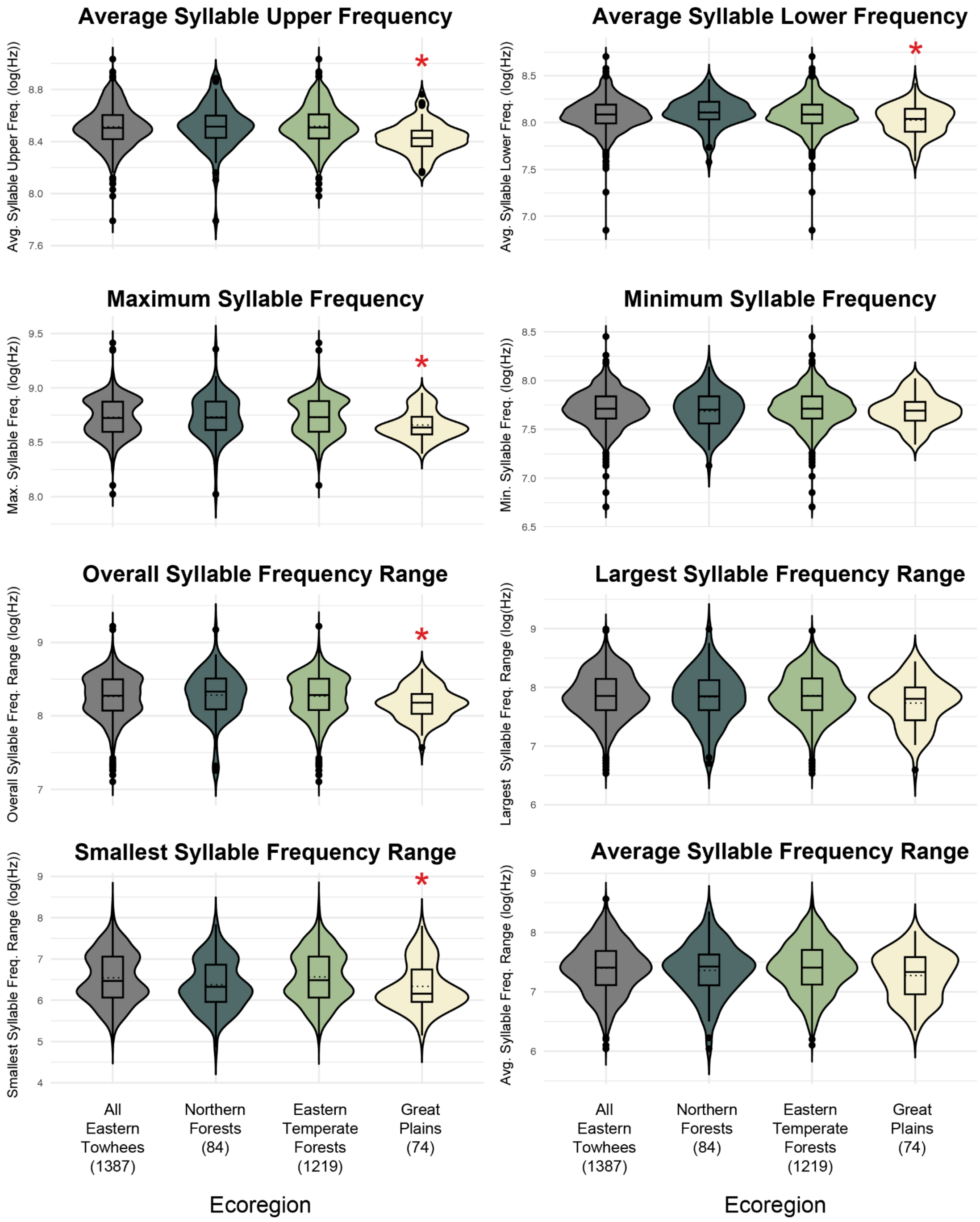


**Supplementary Figure S16.** Violin plots showing the distribution of song frequency traits in each ecoregion for the Eastern towhee in the modern towhee dataset. We used a Wilcoxon rank-sum test to assess if there is a statistically significant difference in song frequency traits in each ecoregion relative to the overall species distribution (N = 1387). Left-most violin on each plot ("All Eastern Towhees") indicates the overall species distribution. An asterisk (*) indicates $p_{adj} < 0.05$ after adjusting for multiple comparisons using the false discovery rate method.

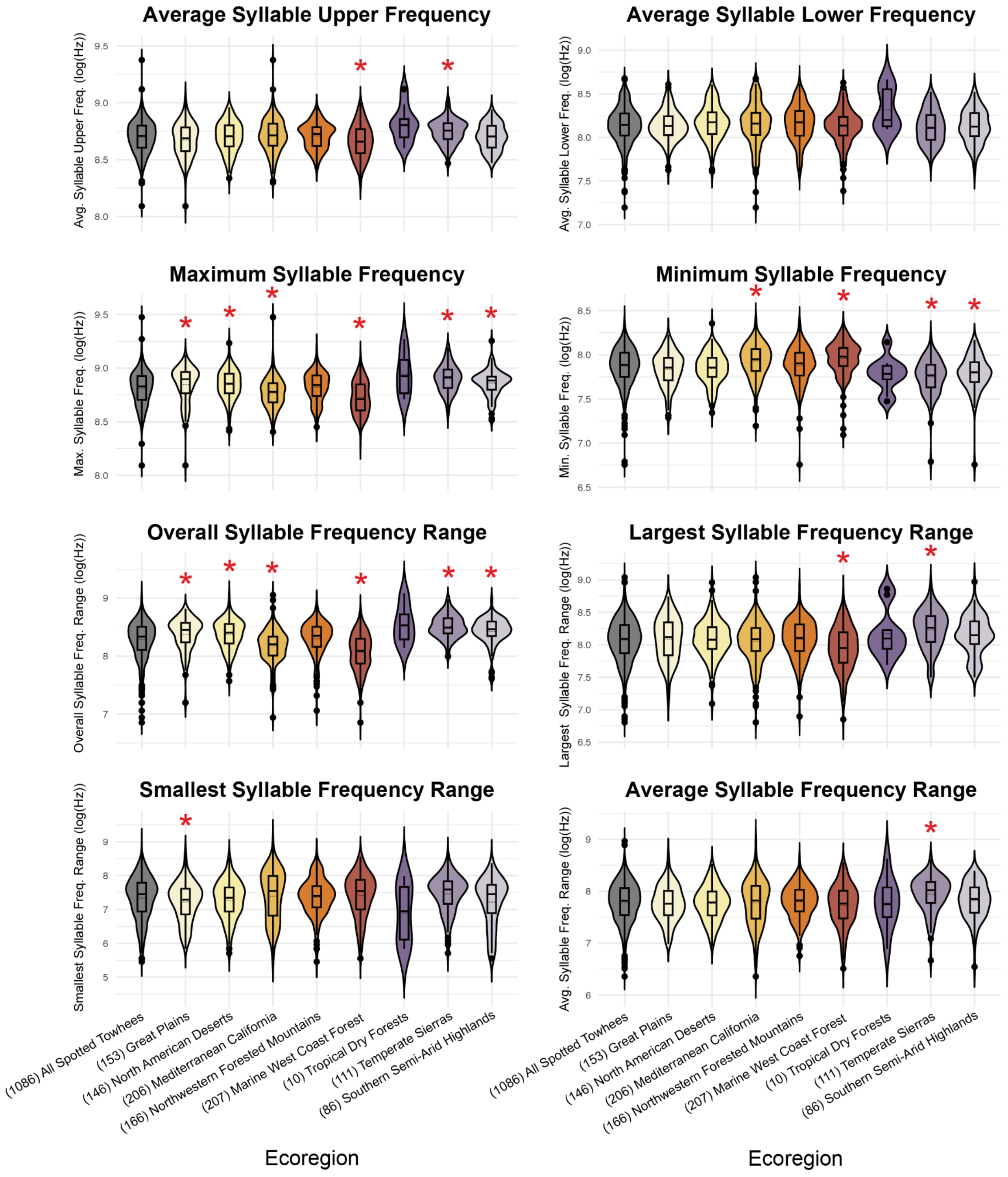


**Supplementary Figure S17.** Violin plots showing the distribution of song frequency traits in each ecoregion for the Spotted towhee in the full towhee dataset. We used a Wilcoxon rank-sum test to assess if there is a statistically significant difference in song frequency traits in each ecoregion relative to the overall species distribution (N = 1086). Left-most violin on each plot

("All Spotted Towhees") indicates the overall species distribution. An asterisk (*) indicates $p_{adj} < 0.05$ after adjusting for multiple comparisons using the false discovery rate method.

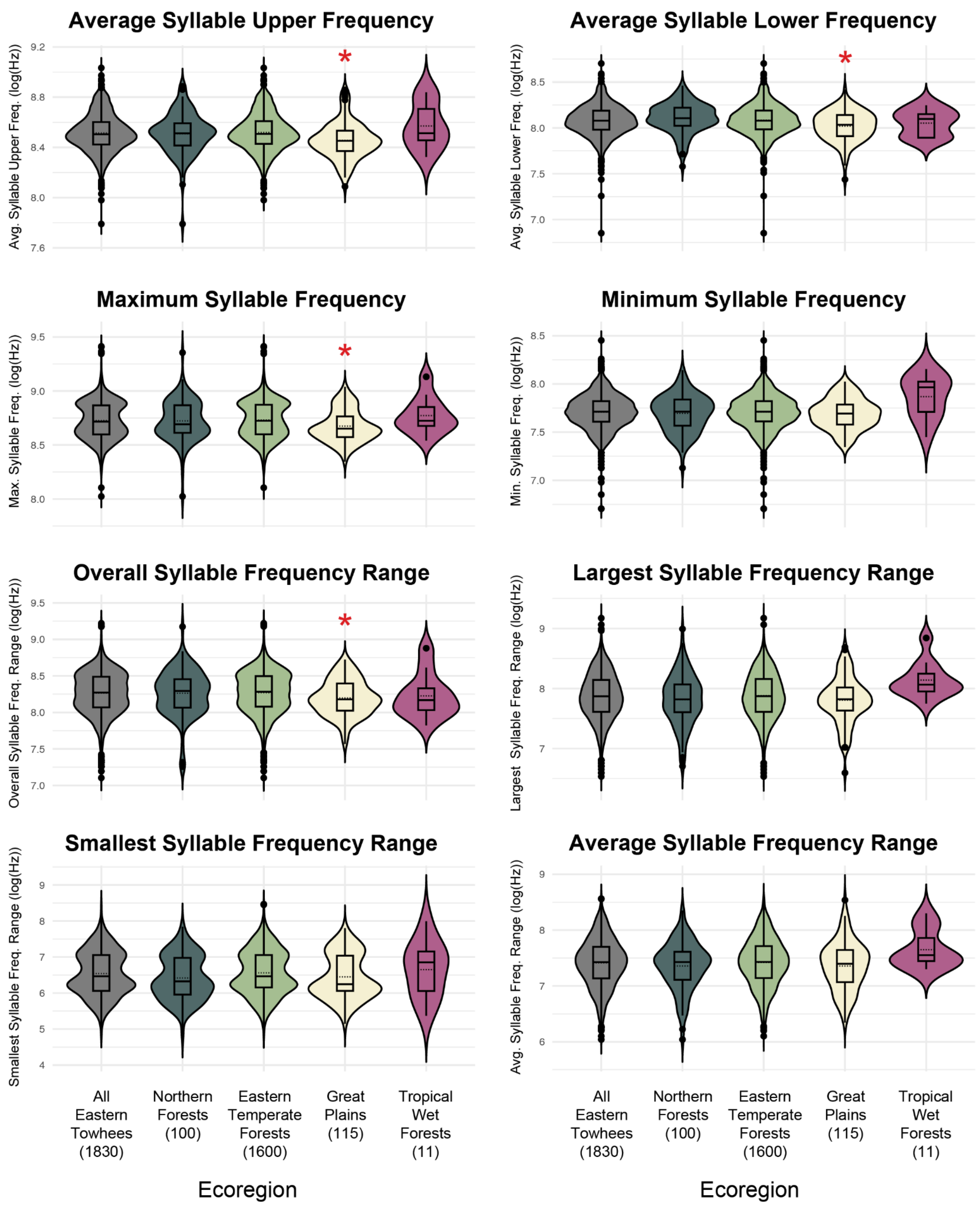


**Supplementary Figure S18.** Violin plots showing the distribution of song frequency traits in each ecoregion for the Eastern towhee in the full towhee dataset. We used a Wilcoxon rank-sum test to assess if there is a statistically significant difference in song frequency traits in each ecoregion relative to the overall species distribution (N = 1830). Left-most violin on each plot ("All Eastern Towhees") indicates the overall species distribution. An asterisk (*) indicates $p_{adj} < 0.05$ after adjusting for multiple comparisons using the false discovery rate method.

**Supplementary Table S5.** Mantel test results for *Song Distance Models* of comparison of geographic distance matrix and multivariate song feature distance matrices of songs of the Eastern towhee (*Pipilo erythrophthalmus*) and the Spotted towhee (*Pipilo maculatus*) in the modern towhee dataset and full towhee dataset.

| Dataset Analyzed | Analysis | Mantel r | *p* |
|---|---|---|---|
| **Modern Towhee Dataset** | Spotted towhee and Eastern towhee combined<br><br>$N_{total} = 2916$<br>($N_{Spotted} = 1086$ ; $N_{Eastern} = 1830$) | 0.171 | **<0.001** |
| | Spotted towhee ONLY<br>N = 1086 | 0.105 | **<0.001** |
| | Eastern towhee ONLY<br>N = 1830 | -0.002 | 0.547 |
| **Full Towhee Dataset** | Spotted towhee and Eastern towhee combined<br><br>$N_{total} = 2916$<br>($N_{Spotted} = 1086$ ; $N_{Eastern} = 1830$) | 0.170 | **<0.001** |
| | Spotted towhee ONLY<br>N = 1086 | 0.121 | **<0.001** |
| | Eastern towhee ONLY<br>N = 1830 | 0.005 | 0.332 |

Results with $p < 0.05$ are bolded and highlighted in red.

**Supplementary Table S6.** Mantel test results of *Song Feature Distance Models* comparing song feature matrices against geographic distance matrices of songs of Eastern towhee (*Pipilo erythrophthalmus*) and the Spotted towhee (*Pipilo maculatus*) in the modern towhee dataset.

| Analysis | Frequency Variable | Mantel r | $p_{adj}$ |
|---|---|---|---|
| Spotted towhee and Eastern towhee combined<br><br>$N_{total}$ = 2098<br>($N_{Spotted}$ = 711 ; $N_{Eastern}$ = 1387) | Average Syllable Upper Frequency (Hz) | 0.166 | **0.002** |
| | Average Syllable Lower Frequency (Hz) | 0.071 | **0.002** |
| | Maximum Syllable Frequency (Hz) | -0.008 | 0.885 |
| | Minimum Syllable Frequency (Hz) | 0.156 | **0.002** |
| | Overall Syllable Frequency Range (Hz) | 0.024 | **0.007** |
| | Largest Syllable Frequency Range (Hz) | 0.009 | 0.118 |
| | Smallest Syllable Frequency Range (Hz) | 0.219 | **0.002** |
| | Average Syllable Frequency Range (Hz) | 0.099 | **0.002** |
| Spotted towhee ONLY<br>N = 711 | Average Syllable Upper Frequency (Hz) | 0.044 | **0.002** |
| | Average Syllable Lower Frequency (Hz) | 0.014 | 0.094 |
| | Maximum Syllable Frequency (Hz) | 0.126 | **0.002** |
| | Minimum Syllable Frequency (Hz) | 0.060 | **0.002** |
| | Overall Syllable Frequency Range (Hz) | 0.151 | **0.002** |
| | Largest Syllable Frequency Range (Hz) | 0.030 | **0.005** |
| | Smallest Syllable Frequency Range (Hz) | 0.029 | **0.005** |
| | Average Syllable Frequency Range (Hz) | 0.015 | 0.093 |
| Eastern towhee ONLY<br>N = 1387 | Average Syllable Upper Frequency (Hz) | 0.016 | 0.752 |
| | Average Syllable Lower Frequency (Hz) | -0.008 | 0.825 |
| | Maximum Syllable Frequency (Hz) | $6.924 \times 10^{-4}$ | 0.825 |
| | Minimum Syllable Frequency (Hz) | 0.004 | 0.825 |
| | Overall Syllable Frequency Range (Hz) | -0.001 | 0.825 |
| | Largest Syllable Frequency Range (Hz) | -0.005 | 0.825 |
| | Smallest Syllable Frequency Range (Hz) | 0.002 | 0.825 |
| | Average Syllable Frequency Range (Hz) | -0.014 | 0.898 |

After FDR correction for multiple comparisons, results with $p_{adj}$ < 0.05 are bolded and highlighted in red.

**Supplementary Table S7.** Mantel test results of *Song Feature Distance Models* comparing song feature matrices against geographic distance matrices of songs of Eastern towhee (*Pipilo erythrophthalmus*) and the Spotted towhee (*Pipilo maculatus*) in the full towhee dataset.

| Analysis | Frequency Variable | Mantel r | $p_{adj}$ |
|---|---|---|---|
| Spotted towhee and Eastern towhee combined<br><br>$N_{total}$ = 2916<br>($N_{Spotted}$ = 1086 ; $N_{Eastern}$ = 1830) | Average Syllable Upper Frequency (Hz) | 0.162 | **0.001** |
| | Average Syllable Lower Frequency (Hz) | 0.068 | **0.001** |
| | Maximum Syllable Frequency (Hz) | 0.002 | 0.324 |
| | Minimum Syllable Frequency (Hz) | 0.159 | **0.001** |
| | Overall Syllable Frequency Range (Hz) | 0.037 | **0.001** |
| | Largest Syllable Frequency Range (Hz) | 0.008 | 0.123 |
| | Smallest Syllable Frequency Range (Hz) | 0.209 | **0.001** |
| | Average Syllable Frequency Range (Hz) | 0.084 | **0.001** |
| Spotted towhee ONLY<br>N = 1086 | Average Syllable Upper Frequency (Hz) | 0.037 | **0.008** |
| | Average Syllable Lower Frequency (Hz) | 0.015 | 0.153 |
| | Maximum Syllable Frequency (Hz) | 0.128 | **0.002** |
| | Minimum Syllable Frequency (Hz) | 0.090 | **0.002** |
| | Overall Syllable Frequency Range (Hz) | 0.149 | **0.002** |
| | Largest Syllable Frequency Range (Hz) | 0.072 | **0.002** |
| | Smallest Syllable Frequency Range (Hz) | 0.020 | 0.101 |
| | Average Syllable Frequency Range (Hz) | 0.033 | **0.019** |
| Eastern towhee ONLY<br>N = 1830 | Average Syllable Upper Frequency (Hz) | 0.032 | **0.016** |
| | Average Syllable Lower Frequency (Hz) | -0.010 | 0.813 |
| | Maximum Syllable Frequency (Hz) | 0.001 | 0.813 |
| | Minimum Syllable Frequency (Hz) | $-2.826 \times 10^{-4}$ | 0.813 |
| | Overall Syllable Frequency Range (Hz) | -0.004 | 0.813 |
| | Largest Syllable Frequency Range (Hz) | $-9.453 \times 10^{-4}$ | 0.813 |
| | Smallest Syllable Frequency Range (Hz) | 0.011 | 0.432 |
| | Average Syllable Frequency Range (Hz) | -0.007 | 0.813 |

After FDR correction for multiple comparisons, results with $p_{adj} < 0.05$ are bolded and highlighted in red.

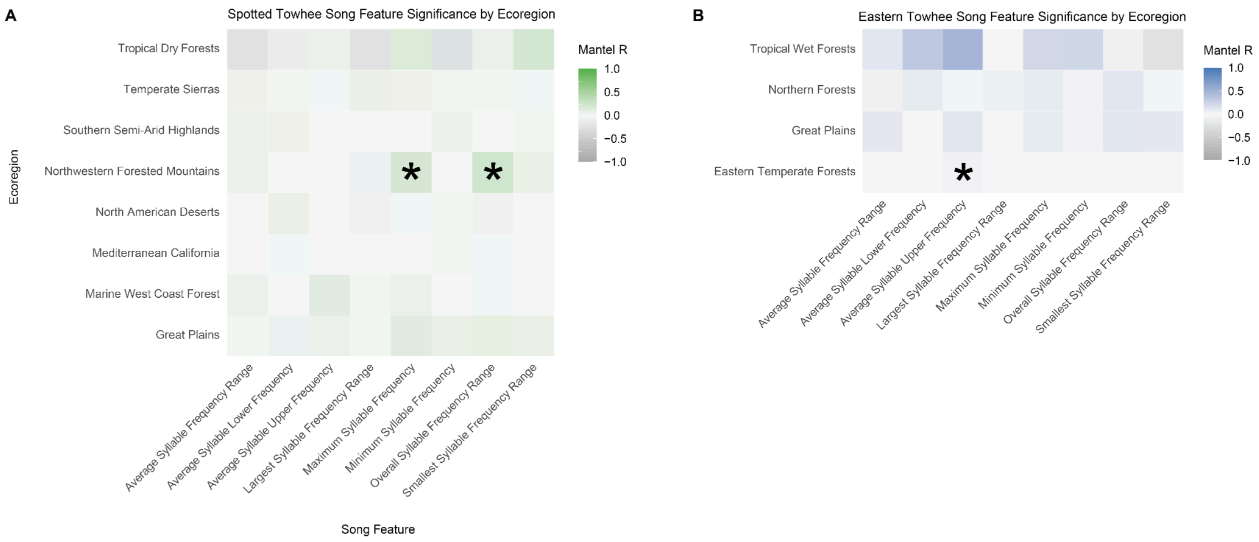


**Supplementary Figure S19.** Heat map of Mantel test testing for evidence for isolation by distance for 8 song features within each ecoregion for songs of the Spotted towhee (*Pipilo maculatus*) and Eastern towhee (*Pipilo erythrophthalmus*) in the full towhee dataset. We computed geographic versus song distance matrices for each ecoregion and song feature pair separately and applied a Mantel test to assess for correlation between geographic distance and song feature distance. An asterisk (*) indicates $p_{adj} < 0.05$ after an FDR correction for multiple comparisons.

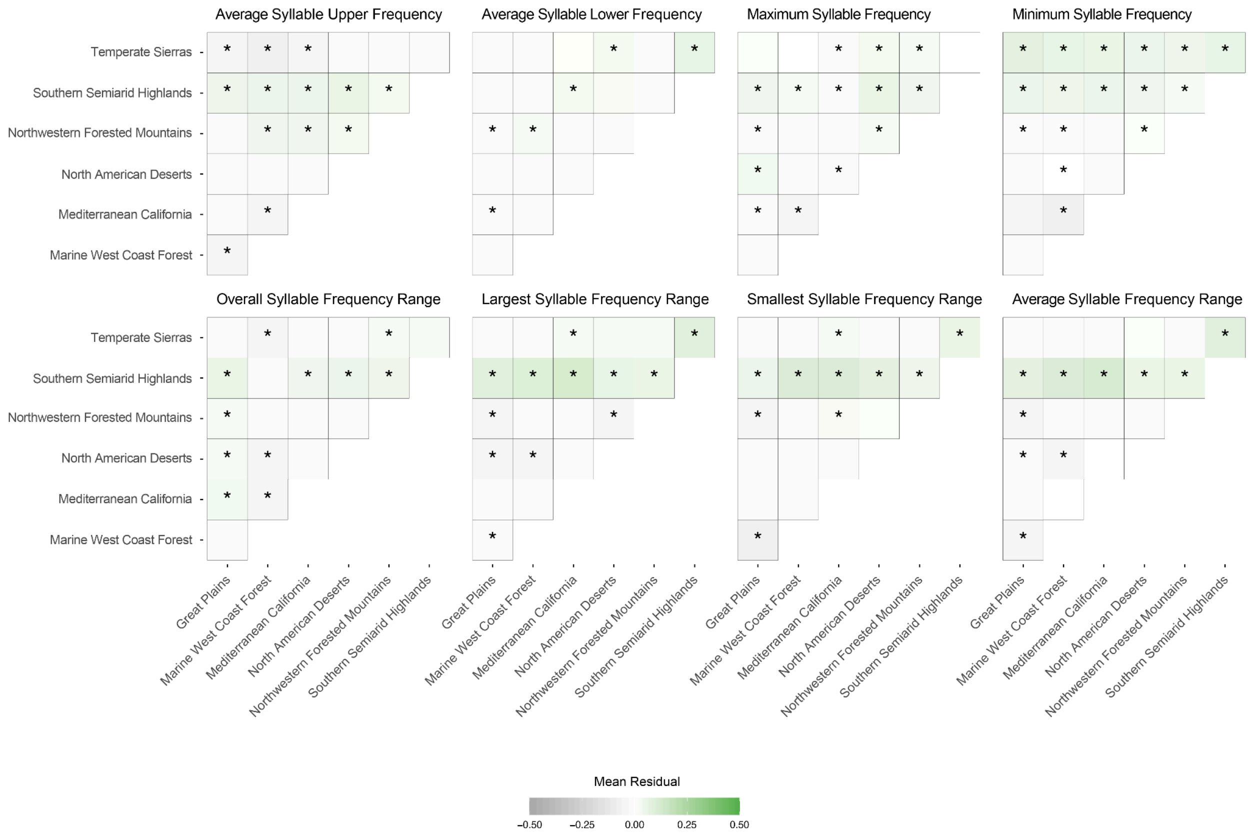


**Supplementary Figure S20.** Heat map describing the mean residual dissimilarity between ecoregions for 8 song features in songs of the Spotted towhee (*Pipilo maculatus*) in the modern towhee dataset. We computed geographic vs. song distance matrices for each possible pair of Spotted towhee samples, and we fit a linear model. We applied a t-test comparing the distribution of residuals between ecoregions against the distribution of residuals within ecoregions. An asterisk (*) indicates $p_{adj} < 0.05$ after an FDR correction for multiple comparisons.

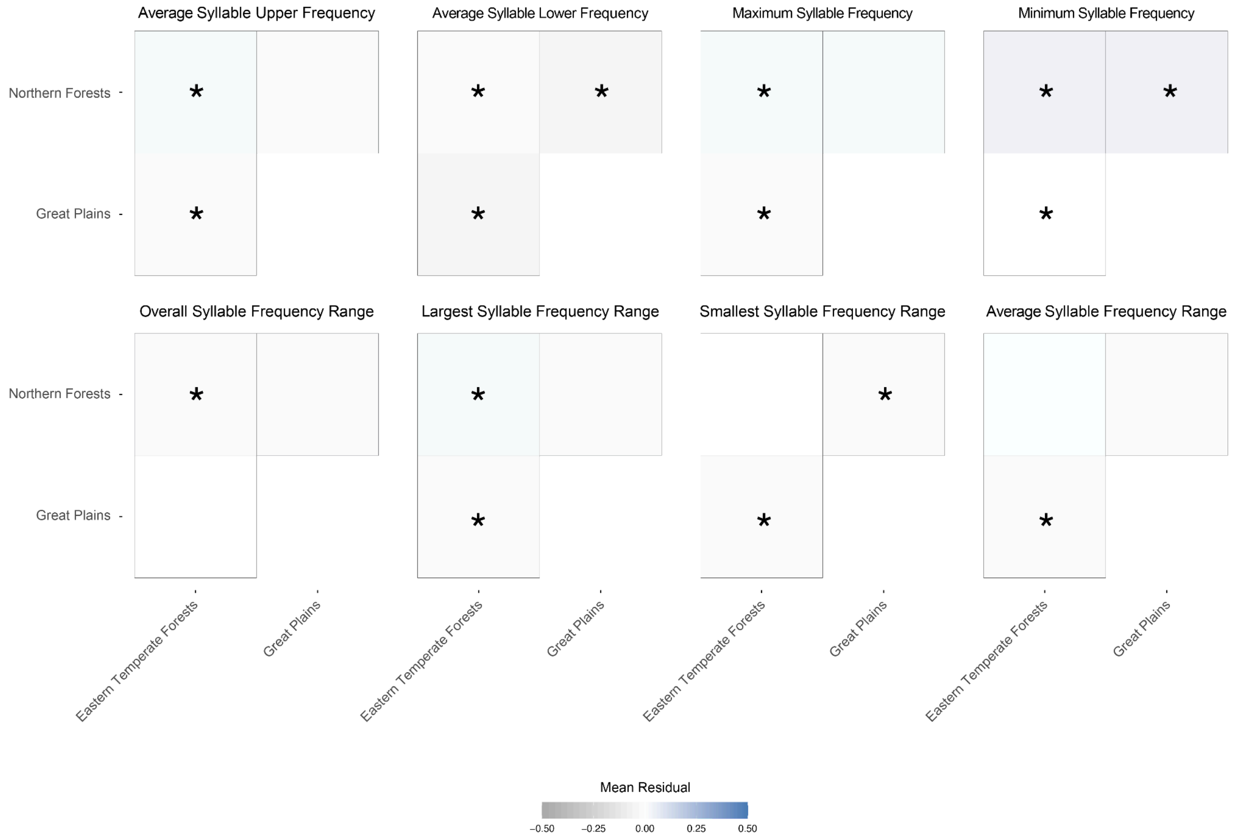


**Supplementary Figure S21.** Heat map describing the mean residual dissimilarity between ecoregions for 8 song features in songs of the Eastern towhee (*Pipilo erythrophthalmus*) in the modern towhee dataset. We computed geographic vs song distance matrices for each possible pair of Eastern towhee samples, and we fit a linear model. We applied a T-test comparing the distribution of residuals between ecoregions against the distribution of residuals within ecoregions. An asterisk (*) indicates $p_{adj} < 0.05$ after an FDR correction for multiple comparisons.

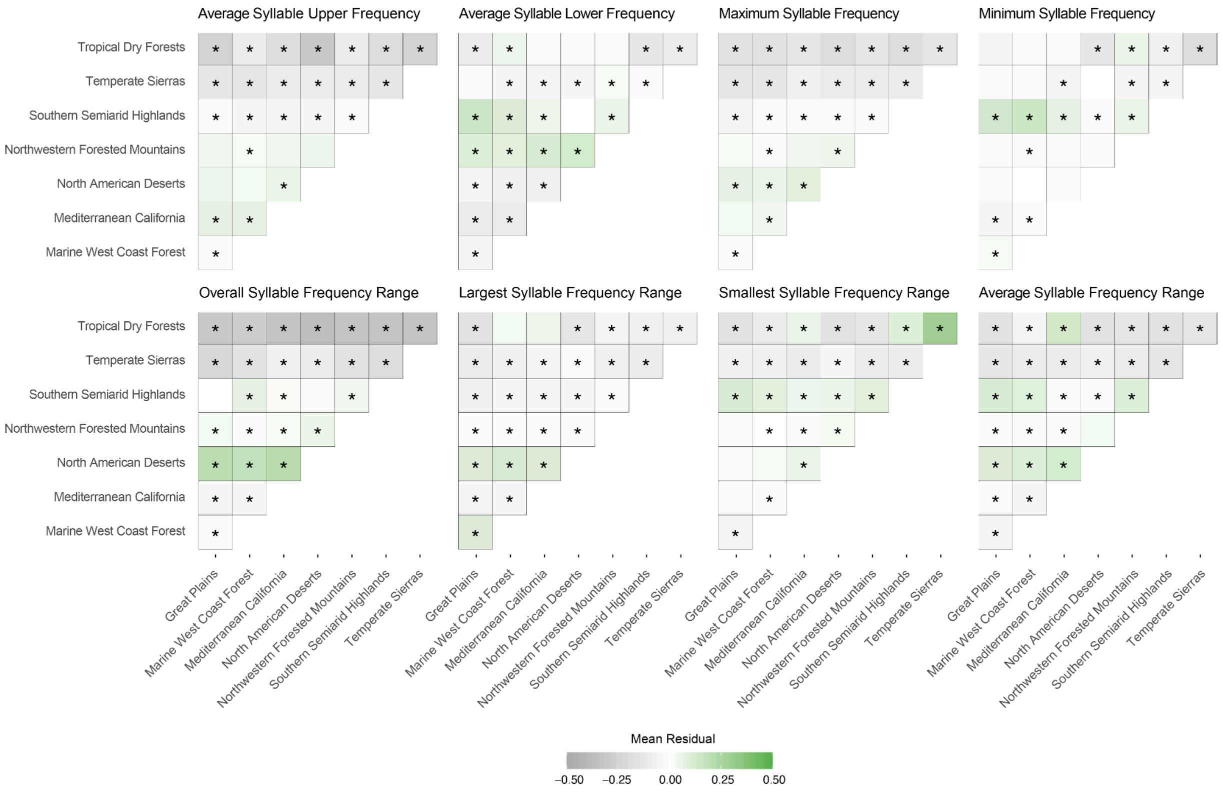


**Supplementary Figure S22.** Heat map describing the mean residual dissimilarity between ecoregions for 8 song features in songs of the Spotted towhee (*Pipilo maculatus*) in the full towhee dataset. We computed geographic vs. song distance matrices for each possible pair of Spotted towhee samples, and we fit a linear model. We applied a t-test comparing the distribution of residuals between ecoregions against the distribution of residuals within ecoregions. An asterisk (*) indicates $p_{adj} < 0.05$ after an FDR correction for multiple comparisons.

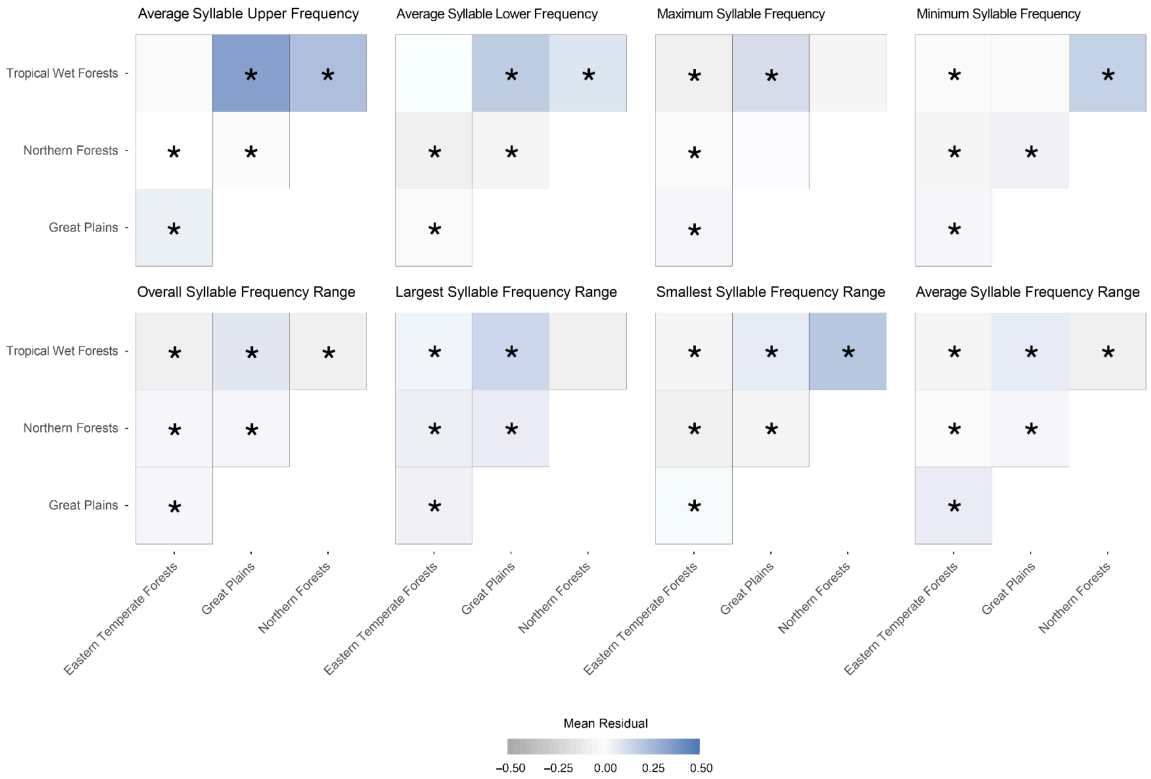


**Supplementary Figure S23.** Heat map describing the mean residual dissimilarity between ecoregions for 8 song features in songs of the Eastern towhee (*Pipilo erythrophthalmus*) in the full towhee dataset. We computed geographic vs song distance matrices for each possible pair of Eastern towhee samples, and we fit a linear model. We applied a T-test comparing the distribution of residuals between ecoregions against the distribution of residuals within ecoregions. An asterisk (*) indicates $p_{\text{adj}} < 0.05$ after an FDR correction for multiple comparisons.
